# Groundwater bacterial communities evolve over time, exhibiting oscillating similarity patterns in response to recharge

**DOI:** 10.1101/2021.03.18.434713

**Authors:** Lijuan Yan, Syrie M. Hermans, Kai Uwe Totsche, Robert Lehmann, Martina Herrmann, Kirsten Küsel

## Abstract

Time series analyses are a crucial tool for uncovering the patterns and processes shaping microbial communities and their functions, especially in aquatic ecosystems. Subsurface aquatic environments are perceived to be more stable than oceans and lakes, due to the lack of sunlight, the absence of photosysnthetically-driven primary production, low temperature variations, and oligotrophic conditions. However, periodic groundwater recharge should affect the structure and succession of groundwater microbiomes. To disentangle the long-term temporal changes in groundwater bacterial communities of shallow fractured bedrock community, and identify the drivers of the observed patterns, we analysed bacterial 16S rRNA gene sequencing data for samples collected monthly from three groundwater wells over a six-year period (n=230) along a hillslope recharge area. We show that the bacterial communities in the groundwater of limestone-mudstone alternations were not stable over time and showed oscillating dissimilarity patterns which corresponded to periods of groundwater recharge. Further, we observed an increase in dissimilarity over time (generalized additive model *P* < 0.001) indicating that the successive recharge events result in communities that are increasingly more dissimilar to the initial reference time point. The sampling period was able to explain up to 29.5% of the variability in bacterial community composition and the impact of recharge events on the groundwater microbiome was linked to the strength of the recharge and local environmental selection. Many groundwater bacteria originated from the recharge-related sources (mean = 66.5%, SD = 15.1%) and specific bacterial taxa were identified as being either enriched or repressed during recharge events. Overall, similar to surface aquatic environments, groundwater microbiomes vary through time, though we revealed groundwater recharges as unique driving factors for these patterns. The high temporal resolution employed here highlights the complex dynamics of bacterial communities in groundwater and demonstrated that successive shocks disturb the bacterial communities, leading to decreased similarity to the initial state over time.

## Introduction

Longitudinal observations of microbial communities can uncover important ecological patterns and processes such as community succession, community stability and responses to environmental disturbances and changes (Faust et al., 2015). Such studies have been used to show the development of the human gut microbiome in an infant (Koenig et al., 2011; Bäckhed et al., 2015), the patterns of plant microbiomes over growing seasons (Redford and Fierer, 2009; Deyett and Rolshausen, 2019) and the stability of bacterial communities in wastewater over the course of a year (Werner et al., 2011). Investigating bacterial communities over long time scales and with high temporal resolution, is a crucial requirement to comprehensively understand microbial community dynamics, their drivers, and their effects on ecosystem processes.

Arguably, the most comprehensive temporal research of environmental microbiomes has been conducted in aquatic systems (Shade et al., 2013). There is evidence of seasonal patterns and annual periodicity in the composition of bacterial communities in lakes (Shade et al., 2007, 2013; Li et al., 2015) and oceans (Fuhrman et al., 2006; Gilbert et al., 2012; Morán et al., 2015; Ward et al., 2017; Wang et al., 2020). Specific marine taxa show preferences to either winter or summer seasons (Ward et al., 2017) and the diversity of the marine bacterial communities is often found to be highest in winter months (Gilbert et al., 2012; Ladau et al., 2013), although this does not always hold true (Ward et al., 2017). However, planktonic bacterial communities in streams do not exhibit seasonal patterns (Portillo et al., 2012), indicating that temporal variation might be system specific, and that not all aquatic bacterial communities show seasonal response patterns.

Unlike limnic and marine systems, groundwater microbiomes have been largely ignored in temporal studies. Subsurface waters are not driven by seasonally varying primary production due to the lack of sunlight, there is little temperature variation, presumably little to no input of surface-derived fresh, easily available organic carbon (Akob and Küsel, 2011; Benk et al., 2019), and groundwater residence time can exceed those of rivers and lakes. It was therefore assumed that groundwater microbial communities would be relatively stable over time (Griebler and Lueders, 2009). However, these systems appear to be less constant than first assumed and there is evidence from short-term studies (< 1 year) that groundwater microbiomes exhibit seasonal dynamics associated with changes in groundwater chemistry and recharge events (Lin et al., 2012; Zelaya et al., 2019; Zhou et al., 2012). It is likely that surface connectivity of an aquifer impacts the amount of temporal variation observed (Küsel et al., 2016); surface-near groundwater especially could harbour temporally distinct microbial communities affecting its ecosystem functioning. We have previously reported that space but not time plays a significant role in shaping bacterial communities when comparing samples collected three months apart over three years but the high spatial variations observed could have masked temporal variabilities (Yan et al., 2020). Finer and longer time scales similar to marine time-series research are needed to identify potential patterns shaping subsurface aquatic systems.

Knowledge of the temporal dynamics of the groundwater microbiome may hold great promise for understanding and predicting the microbial structural and functional responses to precipitation changes and global warming. Stress on karst water resources, which contribute to the drinking water supply for almost a quarter of the world’s population (Ford and Williams, 2007), especially is likely to increase in the future in terms of both water quantity and quality (Hartmann et al., 2014). Therefore, we aimed to disentangle the long-term temporal changes in surface-near groundwater bacterial communities and identify the drivers of the observed patterns. For this study, we took advantage of regular monthly sampling campaigns of groundwater from the monitoring transect of the Hainich Critical Zone Exploratory (CZE; Küsel et al., 2016) in Central Germany. The selected three monitoring wells represent contrasting depths, surface connectivity, and hydrochemistry. We hypothesise that (1) the long observation time (> 6 years) and high temporal resolution of our sampling will allow us to detect temporal variations within the groundwater microbiome, (2) temporal patterns will vary between wells due to varying local conditions and environmental selection strength, and (3) successive disturbances, in the form of groundwater recharge events, will drive the temporal variation through their impact on microbial diversity and abundance, including the introduction of surface-derived taxa.

## Methods

### Study site

In the Hainich CZE, a multi-storey fractured aquifer system within Triassic limestone-mudstone alternations (Kohlhepp et al., 2017) is studied, representing shallow groundwater resources in fractured bedrock. At the Hainich low-mountain ranges’ eastern hillslope (groundwater recharge area), a well transect spans different land use, relief positions and depths (Küsel et al. 2016). Owing to the differences in bulk rock permeability and surface connectivity (Lazar et al. 2019, Lehmann & Totsche, 2020), the laying and limestone-dominated main aquifer (Upper Muschelkalk, Trochitenkalk formation) is well-oxygenated whereas the hanging and mudstone-dominated aquifer assemblages (Meissner formation) are partly oxygen-deficient, contributing to distinct biogeochemical zones with varying microbial community assembly mechanisms and metabolic processes (Wegner et al., 2018; Yan et al., 2020).

Three wells, representing different biogeochemical conditions, were selected for the analysis of long-term microbial time-series. Well H41 (48 m below ground level) is located at the lower midslope and accesses the (lower) main aquifer, whereas well H43 (same location, 12 m bgl), and well H52 (65 m bgl), at the Hainich’s footslope, access the upper aquifer assemblage (Yan et al., 2020; Figure 1A). In situ monitoring and quality fluctuations, including groundwater temperatures, are described in Lehmann & Totsche (2020). Cumulative precipitation varied between years and was low (< 600 mm) in 2013, 2015, 2016 and 2018, but highest (860 mm) in 2017 (Figure 1B).

**Figure 1.**
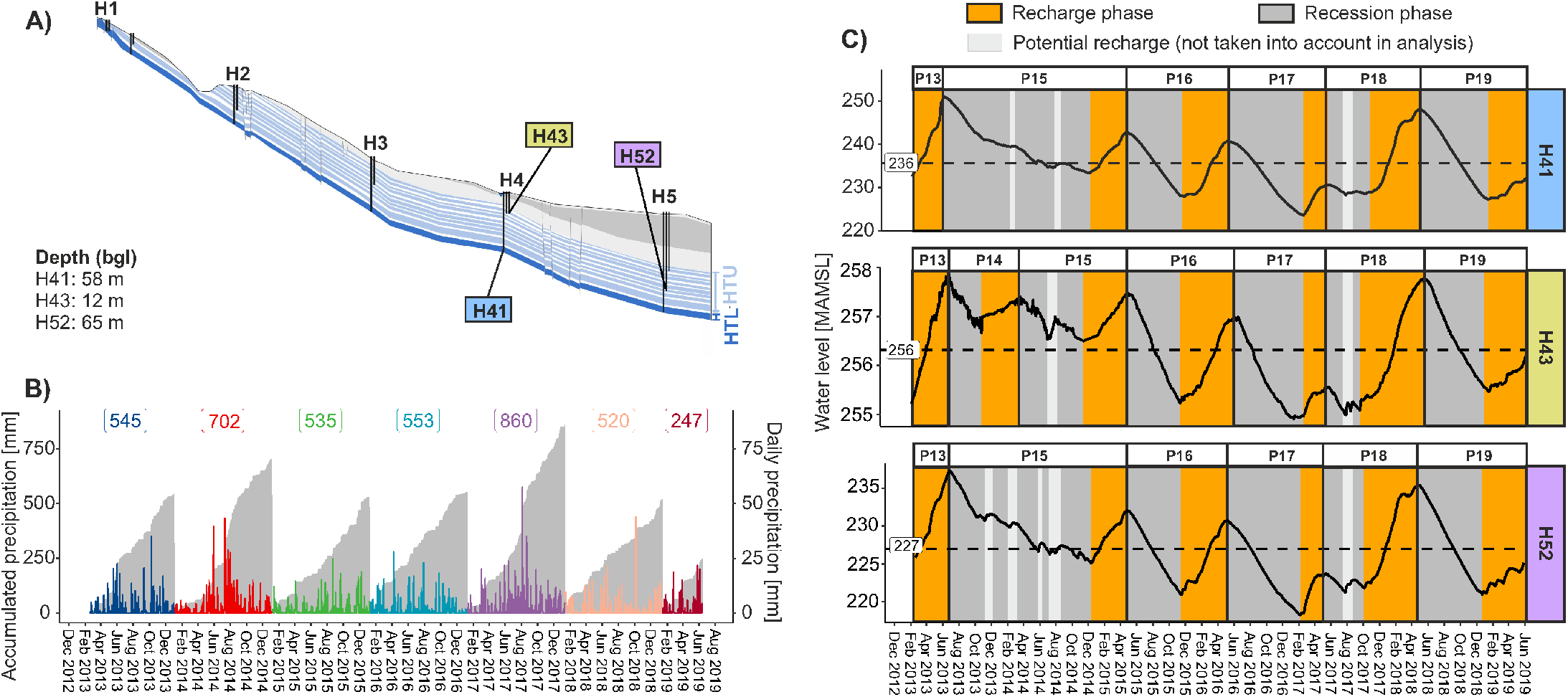
Transect information: A) location of the three wells along a single hillslope of the Hainich transect, modified from Kohlhepp et al. (2017), B) daily (peaks) and accumulated (grey shaded area) precipitation during the investigation period, C) dynamic changes of groundwater level in different wells on a daily basis. The mean water level of the investigation period for each well was indicated with a value box and a dashed line. “Recharge phases” were defined for hypothesis testing based on water level changes (increase: recharge; decrease: recession). A period, labelled as “P + two-digit year”, included one recharge phase and one recession phase, except for P13 which only had a recharge phase. Abbreviations: bgl depth in meters below ground level (m), MAMSL meters above mean sea level.

### Sampling

Groundwater samples were collected on a 4-week basis using submersible sampling pumps (MP1, Grundfos), as previously described (Küsel et al. 2016, Lehmann & Totsche, 2020) from February 2013 to May 2019, allowing us to explore the temporal dynamics of the bacterial communities and hydrochemistry over 75 months. In total, 230 samples were collected over the course of the study. During each sampling campaign, 5-10 L groundwater per well were collected for microbiological analyses in sterilized bottles, after physicochemical parameters stabilized. The determination of hydrochemical parameters was performed as described earlier (Kohlhepp et al., 2017).

### DNA extraction, quantitative PCR, and amplicon sequencing

Genomic DNA was immediately enriched on conventional-sized (0.2 μm) PES (Supor, Pall Corporation, until June 2016) or polycarbonate filters (Nuclepore, Whatman; Merck-Millipore, after June 2016) and kept at −80 °C until DNA extraction. Genomic DNA was subsequently extracted using the PowerSoil DNA Isolation kit (MO BIO Laboratories Inc., USA) and kept at −20 °C until PCR amplification.

Quantitative PCR was performed on a Mx3000P instrument (Agilent Technologies) using Brilliant II SYBR Green qPCR Mastermix (Agilent Technologies) and the primers Bac8Fmod (5’-AGA GTT TGA TYM TGG CTC AG-3’) and Bac338Rabc (5’-GCW GCC WCC CGT AGG WGT-3’) (Daims et al., 1999; Loy et al., 2002) for bacterial 16S rRNA gene following cycling conditions previously reported (Herrmann et al., 2012). Three technical replicates per sample were used to increase the reproducibility (coefficient of variation < 15%).

PCR amplification of bacterial partial 16S rRNA gene (V3-V4 region) was performed using the primers Bakt_0341F (5’-CCTACGGGNGGCWGCAG-3’) and Bakt_0785R (5’-GACTACHVGGGTATCTAATCC-3’) (Herlemann et al., 2011; Klindworth et al., 2013) according to the PCR programs described by Klindworth et al., (2013). The amplicons were sequenced on Illumina’s MiSeq platform using V3 (2 x 300 bp) chemistry. Most samples (as listed in Supplementary Table S1) were sequenced in-house using library preparation procedures as described by (Kumar et al., 2018). The remaining samples (as listed in Supplementary Table S1) were sequenced at LGC genomics GmbH (Berlin, Germany), following similar methods as for the in-house samples (Rughöft et al. 2016).

### Bioinformatics analysis

Sequence analysis was performed using mothur V.1.39.5 (Schloss et al. 2009), following the MiSeq SOP (Kozich et al., 2013). Sequences containing ambiguous (N) bases or homopolymers longer than 8 nucleotides were removed. The remaining sequences were pre-clustered, allowing up to 1 bp difference per 100 bp, to remove potential sequencing errors prior to the identification of the chimeric sequences using the UCHIME algorithm (Edgar et al., 2011). After removing chimeric sequences, the sequences were classified against the mothur-formatted SILVA v. 132 taxonomy reference database (Quast et al., 2013), using the default bootstrapping algorithm (cutoff value: 80%). Only sequences classified as Bacteria were retained and clustered into OTUs based on VSEARCH abundance-based greedy clustering (AGC) method at 97% similarity (Rognes et al., 2016). The raw sequencing data were deposited in the European Sequence Archive (ENA), with BioProject IDs and accession numbers as shown in Supplementary Table S1.

### Statistical analysis

Statistical analyses were done using R version 3.5.2 and we used a significance level of α = 0.05 for all statistical tests. All R scripts for the statistical analyses are available at: https://github.com/pikatech/cyclic_patterns.

Periods of increased seasonal recharge were identified in groundwater level time series as the periods between annual minima and maxima. Recession periods represent falling water levels, generally due to limited surface-recharge. In general, a period, here labelled as “P + two-digit year”, included one recession phase and one recharge phase. P13 however, only consists of a recharge phase.

The OTUs with relative abundance less than 0.005% were considered as run-to-run PCR and sequencing errors (Bokulich et al., 2013) and were therefore removed. To fix the variation in sampling depth (library size), cumulative Sum Scaling (CCS), a median-like quantile normalization method, was performed using R package metagenomeSeq (Paulson et al., 2013), prior to multivariate data analysis.

We plotted bacterial community dissimilarity (Bray-Curtis distance) against sampling time interval (days) for all pairwise observations from the same well and a smooth curve was added using ‘geom_smooth’ to illustrate the non-linear relationship between sampling time difference (days) and community dissimilarity. Generalized additive models (‘gam) were used as the smoothing method. The smooth terms were all significant (*P* < 0.001) for all three wells, indicating clearly nonlinear effect of time on bacterial community dissimilarity.

To visualize dissimilarities of bacterial community composition in each groundwater well, an unconstrained ordination (principal coordinates analysis, PCoA) was performed on the relative abundance data based on Bray-Curtis distance. Variation partitioning analysis was done based on a 2-factorial PERMANOVA model (Bray-Curtis distance ∼ Period * Recharge), with significance tested using the default 999 permutations.

Distance-based redundancy analysis (db-RDA) was performed to test the relationships between the bacterial communities and the standardized hydrochemical parameters in each well based on Bray-Curtis distance with 999 permutations using R package Vegan (Oksanen et al., 2007). Linear dependencies between constraints were investigated via the variance inflation factor VIF. Factors with VIF >10 and non-significant factors (based on 999 permutations) were removed from the model.

The Shannon alpha diversity index was estimated from the raw data (without normalization) using the R package Phyloseq (McMurdie & Holmes 2013). Since the values of the Shannon diversity and bacterial 16S rRNA gene abundance (estimated from quantitative PCR) were not normally distributed (based on Shapiro-Wilk test of normality), we used non-parametric methods for hypothesis testing. Non-parametric one-factorial Kruskal–Wallis test was used to test the difference of bacterial diversity and abundance between the three wells. If there was significant difference between the wells, Dunn test was used for multiple comparisons using R package FSA (Ogle, 2015). The p-values for multiple comparisons were adjusted using the Benjamini-Hochberg method (Benjamini and Hochberg, 1995). The difference in bacterial alpha diversity and abundance between the recharge and the recession phases of each period was tested using non-parametric Wilcoxon method.

To unravel the effect of recharge events on the groundwater bacterial communities in each period, we performed fast expectation-maximization for microbial source tracking FEAST (Shenhav et al., 2019). The FEAST method estimates the contribution (in proportion) of the input source conditions to the sink communities and also reports the fraction of the unknown source contribution (i.e. 100% – the fraction of the known source contribution). Here, we used the recharge samples as sinks and the recession samples (before the corresponding recharge period) as sources in each period. Thus, the contribution of the unknown source calculated could be used to quantify the effect of recharge on the bacterial communities in each period.

Non-parametric Kruskal–Wallis tests were used to test the differences in the proportion of unknown source between the defined periods for each well. If there was significant difference of the unknown source contribution between the periods, Dunn test was used for multiple comparisons using R package FSA (Ogle, 2015). The p-values for multiple comparisons were adjusted using the Benjamini-Hochberg method (Benjamini and Hochberg, 1995).

To determine the bacterial markers that discriminated bacterial communities between phases of increased recharge and recession phases at all taxonomic levels, linear discriminant analysis effect size LEfSe (Segata et al., 2011) was performed (threshold: *P* < 0.05, LDA effect size > 2). Due to the high temporal difference in the bacterial community data, we only aimed to discover the bacterial taxa that were associated with recharge or recession phases, irrespective of the period (P14-P19) (class: recharge; subclass: period). Our dataset did not include a recession phase in P13 for pairwise comparison between recharge and recession phases so P13 data were not considered for LEfSe analysis.

## Results

### Groundwater level and quality fluctuations in the aquifers

The groundwater levels in the fractured, mixed carbonate-siliciclastic bedrock predominantly showed seasonal patterns with highstands in spring/early summer and lowstands in early winter. Maximum water level fluctuations at H41 reached 27.3 m, followed by H52 with 19.0 m and 3 m in the shallow well H43 (Figure 1C; Figure S1, S2, S3). Groundwater recharge usually occurred from October to March/April with peaks owing to snowmelt. During summer recessions, minor rises of water levels reflect groundwater recharge events, predominantly after rainstorms with extreme amounts of precipitation (>99.9th percentile, see Lehmann & Totsche, 2020). We assigned six (H41 and H52) or seven (H43) periods, consisting of a consecutive recession and recharge phase (Figure 1B-C).

In the shallower groundwater (well H43), typically sinusoidal temperature fluctuations occurred, with a mean of 9.2°C and amplitudes of 0.4 K that were delayed by 7 months in comparison to the air temperature. Recharge events only caused minor variation <0.1 K, mostly cooling. At the same location, in a depth of 48 m (mean: 9.2°C), seasonal temperature variation was ∼0.3 K, whereas in the discharge direction, temperatures in 65 m depth (mean: 9.6°C) fluctuated by <0.1K, both coinciding with seasonal water level fluctuation.

The well H41 show considerable fluctuation in Ca, SO_42-_, dissolved oxygen, NO_3-_with fluctuating water levels and inverse variation of K, Mg, Na, Si, and NH_4+_. Contrastingly, the shallow well H43 show insignificant differences between recharge and recession phases only. In comparison to the deep midslope well, the deep well in discharge direction (H52) is supplied by NH_4_^+^ during recharge phases and by NO_3_^-^ during recessions, whereas SO_4_^2-^, Mg, and Na vary correspondingly.

### Groundwater bacterial community composition varied over time

We assessed statistically significant temporal patterns in the dissimilarity of bacterial communities in all three wells. The relative abundances of the most dominant classes varied over time within each well (Figure 2A-C). With increasing time between two compared samples, the dissimilarity of the bacterial communities in those samples also increased in a non-linear manner (Gam *P* < 0.001 for all wells, Figure 2D-F). Bacterial community dissimilarity, as reflected by the median Bray-Curtis distance values, was lowest in the H52 (0.56) and higher in H41 (0.64) and H43 (0.67).

**Figure 2.**
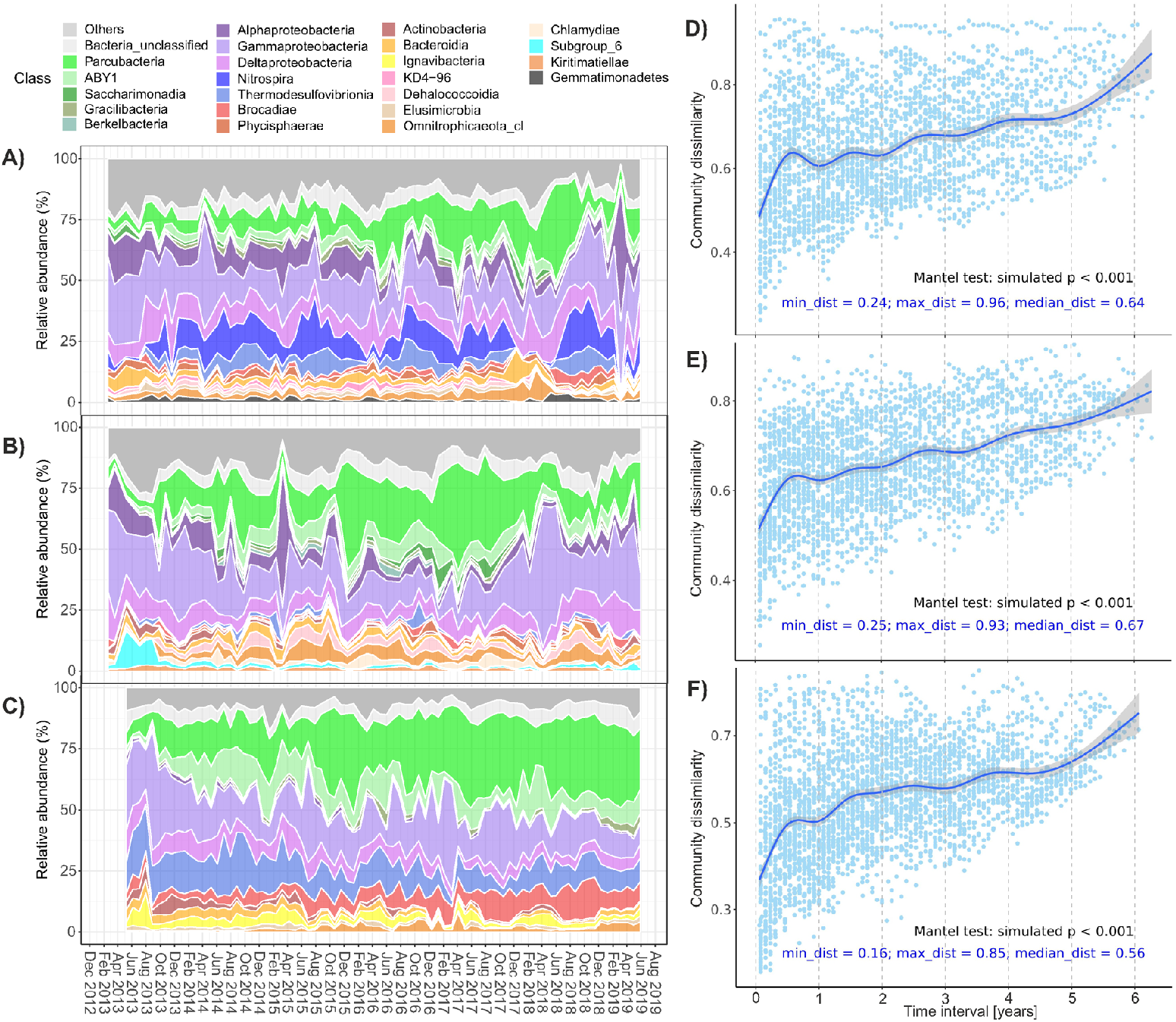
Bacterial community compositional changes over time. Taxonomic composition of A) H41, B) H43 and C) H52 at the class level. The values (mean and standard deviation) of the relative abundance of each class or phylum in the three wells were reported in Supplementary Tables S2-4. Changes in community dissimilarity (Bray-Curtis distance) of D) H41, E) H43 and F) H52 over time. Blue lines represent a smooth curve which was added using “gam” (generalized additive model) as the smoothing method. The smooth terms were significant (P < 0.001) for all three wells. Abbreviations: Others bacterial classes with mean relative abundance less than 1% in the groundwater bacterial community in each well.

Within a well, samples collected at different times of the year, especially when sampling was separated by roughly 6, 18 or 30 months, contained more dissimilar bacterial communities than samples collected at the same time of year (i.e. 12, 24 or 36 months apart; Figure 2D-F). This resulted in an oscillating pattern in bacterial community dissimilarity (Figure 2D-F). This oscillating dissimilarity pattern was weakened over time and when the time interval between two samples was greater than three years the pattern was no longer obvious. After 6 years, the communities exhibited a dissimilarity of over 0.7 in H52 and 0.8 in H41 and H43 (Figure 2D-F).

### Factors relating to changes in bacterial community over time

Differences in groundwater bacterial community structure in all three wells was significantly related to the different periods (P13-P19) and phases (recharge/recession). Period alone explained the largest fraction of the variation in the bacterial communities in all three wells; the explained variation was highest in H52 (29.5%) and lowest in H41 (20.3%) (Figure 3B, D and E). The variation in bacterial community composition explained by the recharge events was dependent on the period and ranged from 17.6% (including single term and the interaction term effects) of variation in the groundwater bacterial communities in H41, to 12.2% in H52. The strong relationship between community composition and phase (recharge/recession) can also be observed along the first PCoA axis for wells H41 (Figure 3A) and H43 (Figure 3C), as the samples show clear separation according to if they came from a recharge or recession phase.

**Figure 3.**
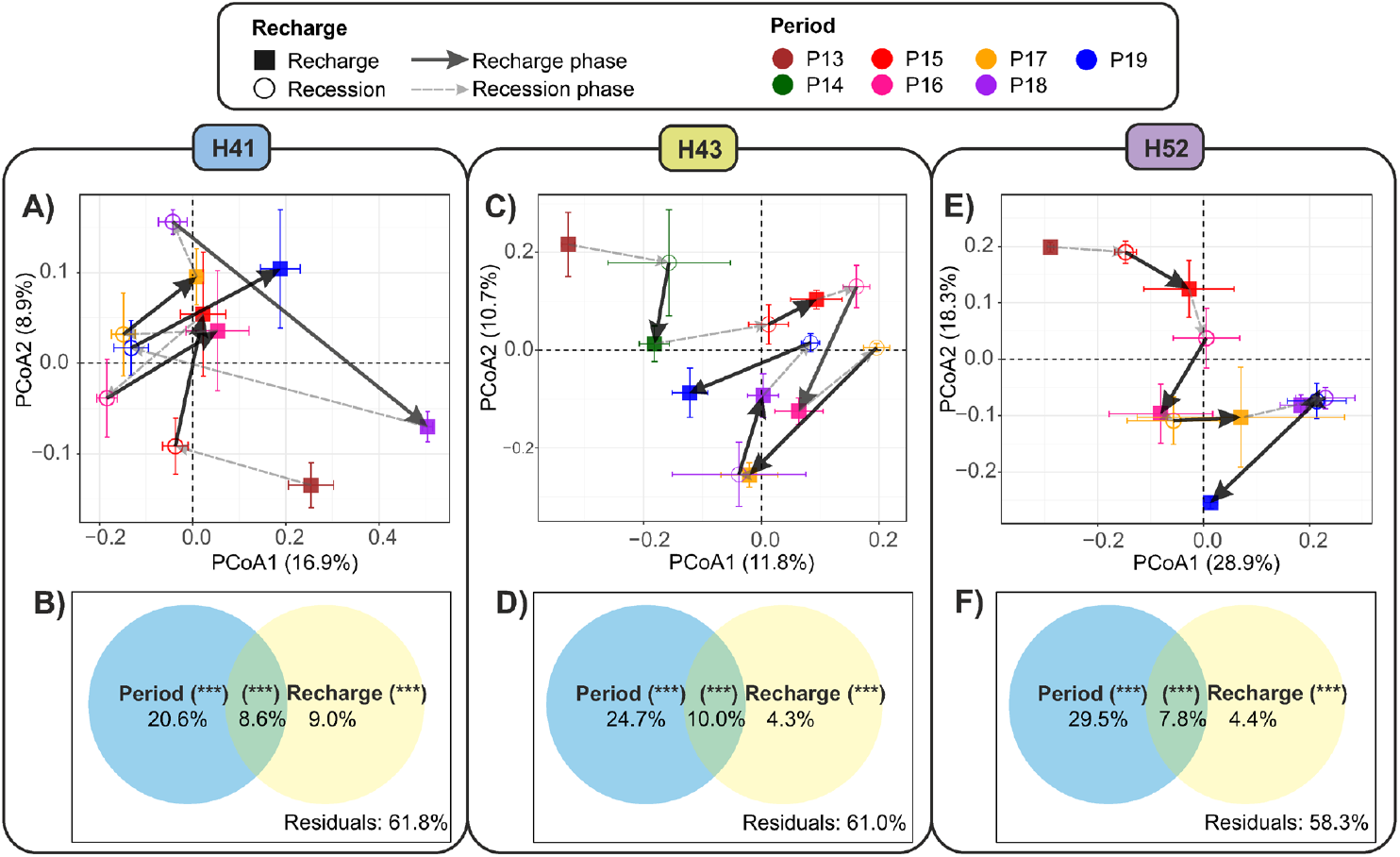
Unconstrained ordinations (PCoA) showing the changes in bacterial community composition between recharge and recession phases and among different time periods in A) H41, C) H43 and E) H52 wells. The points are the centroid of the groups and the error bars represent the standard error of the means of each group in the first two axes of the PCoA ordinations. The effect (trajectory) of recharge (or recession) was shown as the distance (connected with arrows) between centroids of the two successive phases over time. Partitioning of variation across the multivariate bacterial community data was performed based on a 2-factorial PERMANOVA model (Bray-Curtis distance ∼ Period + Recharge + Period:Recharge) for B) H41, D) H43 and F) H52 wells. The significance was computed with 999 permutations (***: p < 0.001).

The effect of recharge on bacterial diversity and abundance was well-and period-dependent. Recharge significantly increased bacterial diversity in H41 (Wilcoxon, p < 0.01). In P18, bacterial diversity in H41 was markedly increased by 14% by recharge. Recharge significantly elevated bacterial abundance in H43 (Wilcoxon, p < 0.05), by 17% in P14 and by 200% in P19, respectively (Supplementary Table S5). However, bacterial diversity was decreased by recharge by 21% in P18 in H43 (Supplementary Table S6). Bacterial diversity and abundance remained unaffected by recharge in H52 over the investigation period (Supplementary Table S5, S6).

Changes in bacterial community composition also correlated with changes in water level, nitrate, and ammonium that fluctuated over time, and explained 16.1%, 9.5% and 15.2% of variations in wells H41, H43 and H52, respectively (Supplementary Figure S4). Contrasting effects were observed in the wells. In H41, changes in bacterial community composition were negatively correlated with nitrate and ammonium during recharge (Supplementary Figure S4A), whereas a positive correlated with dissolved oxygen and nitrate was observed in H43 (Supplementary Figure S4B). In H52, changes in bacterial community composition were positively correlated with nitrate and negatively correlated with ammonium during recharge (Supplementary Figure S4C).

### Specific bacterial taxa can be associated with recharge phases

The contribution of recharge-related taxa to the baseline communities (i.e. those present during recession phases) in the groundwater were well-and period-dependent (Figure 4). Among the three wells, bacterial communities in H43 received the highest taxa contribution from the recharge-related source (mean = 41.0%, SD = 16.5%) over the investigated periods, followed by H41 (mean = 33.5%, SD = 23.3%), and significant differences in the contribution of the recharge-related taxa between periods were evident in these wells. In P18, the overwhelming majority of bacteria originated from the recharge-related source in both H41 (mean = 66.5%, SD = 15.1%) and H43 (mean =57.8%, SD = 15.4%). The contribution of recharge-related taxa was smallest in the periods between P15 and P17 when the water level change during recharge was only minor. The contribution of recharge-related taxa was consistently low (mean = 9.1%, SD = 4.6%) over the periods in well H52.

**Figure 4.**
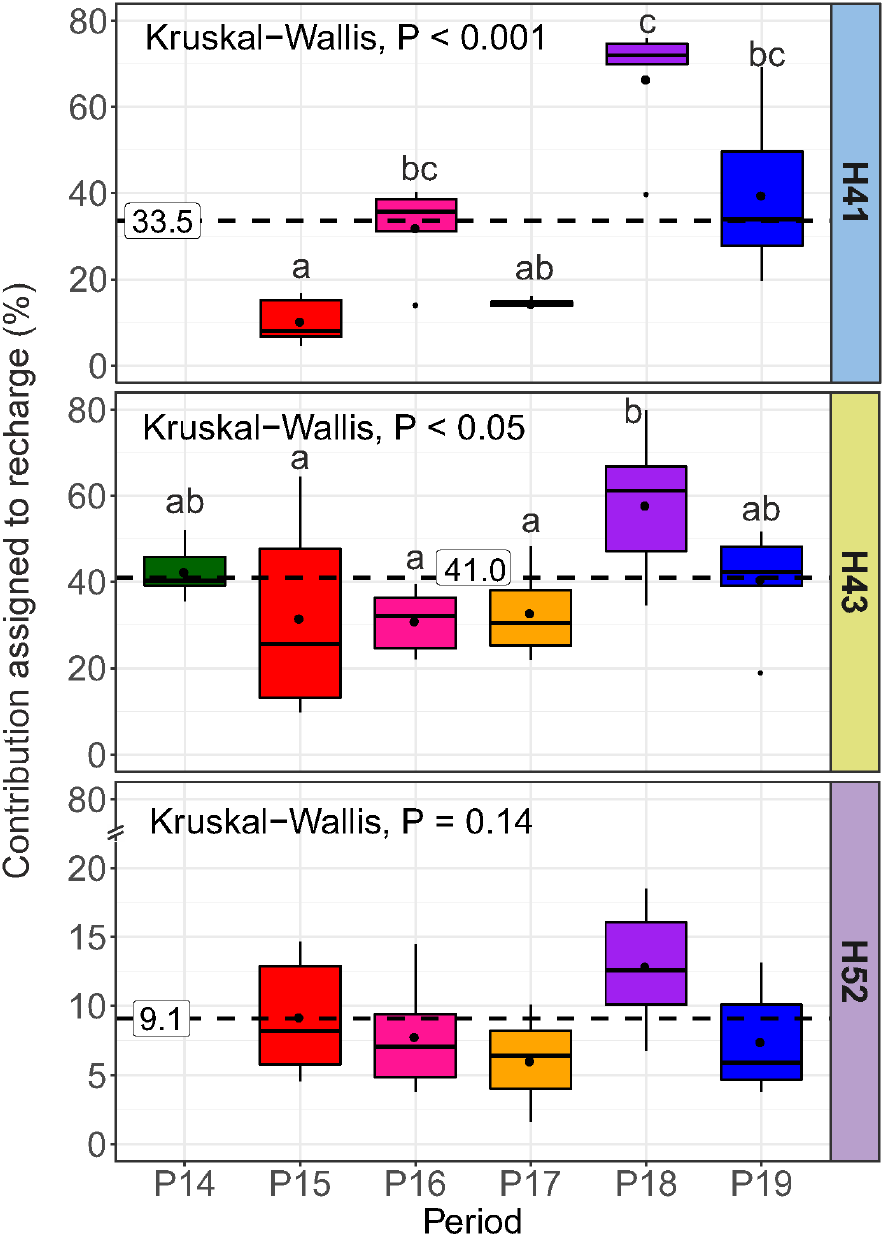
The proportion of taxa contributed to the bacterial communities during recharge in each defined period. Dashed lines and boxed numbers represent the mean contribution across all periods within each well.

We observed considerably more taxa, across all taxonomic levels, that were consistently associated with recharge events in well H41 (231) compared to wells H43 and H52 (56 and 73 respectively; Supplementary Figure S7-9). Well H41 had slightly more recharge favoured taxa (55.4%) than recharge repressed, while in wells H43 and H52 there were more recharge-depressed taxa (89.3% and 54.8%, respectively).

At the genus level, there were 84 discriminative taxa in well H41, of which 47 were recharge favoured (Figure 5A). These genera were predominantly from within the Proteobacteria phylum, although recharge favoured genera also included Patescibacteria such as Saccharimonadales and Ca. Peribacteria. Overall, much fewer discriminative genera were identified in wells H43 and H52 and most were recharge-depressed (89.5% and 69.2% respectively). Nitrospira and Thermodesulfovibrionia were the most recharge-depressed taxa in H41. The recharge-depressed genera in well H43 came from nine different phyla and included several uncharacterised or unclassified genera while most recharge-depressed genera in H52 belonged to the Proteobacteria phylum (Figure 5).

**Figure 5.**
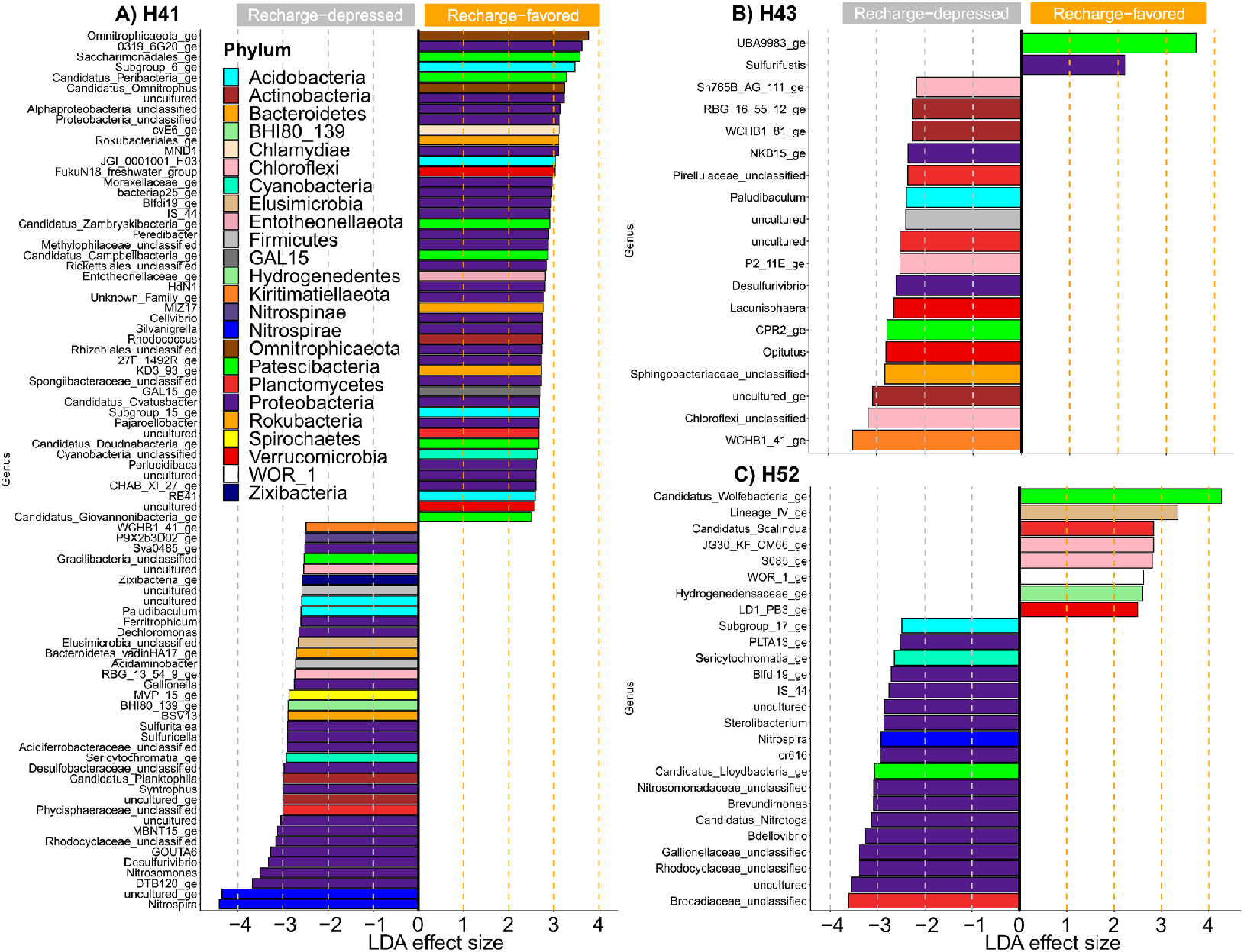
Discriminative features of groundwater bacteria taxa between recharge and recession phases in at the genus level in the three wells, tested using LEfSe (LDA > 2, P < 0.05). ‘Recharge-depressed’ taxa were under-represented during recharge events, while ‘recharge-favored’ taxa 660 were over-represented.

## Discussion

By analysing a comprehensive time-series of monthly samples collected across three wells in a limestone geological setting, we demonstrated that the groundwater microbiome shows significant alterations. These changes were in response to groundwater recharge in shallow and deeper zones of fractured bedrock aquifers, revealing considerable microbial succession and evolution over the 6-year sampling period. The temporal patterns we observed highlight that the subsurface is much more dynamic than previously thought and stress the need to investigate temporal patterns not only in surface aquatic habitats, but also across a range of characteristics within subsurface aquatic systems.

The strong cyclical patterns observed in surface aquatic microbial communities were brought to light because of the high temporal resolution and long timespans employed by the studies (Shade et al., 2007; Gilbert et al., 2012; Morán et al., 2015); similarly expansive temporal research had been lacking for groundwater ecosystems. Studies conducted over shorter time scales, or less frequent sampling intervals nonetheless pointed to seasonal differences in the groundwater microbiome (Zhou et al., 2012), and fluctuations in the abundances of consistently present taxa (Hubalek et al., 2016; Zelaya et al., 2019). By employing our long-term, high resolution sampling and analysis approach, we demonstrated strong oscillating patterns of subsurface aquatic microbial communities. However, these patterns were weaker and not cyclic like what is observed in marine environments (Fuhrman et al., 2006; Shade et al., 2007, 2013; Gilbert et al., 2012; Li et al., 2015; Morán et al., 2015; Ward et al., 2017; Wang et al., 2020). For marine environments, seasonal patterns are strongly linked to changes in temperature, salinity and the change in day length (Fuhrman et al., 2006; Ward et al., 2017), while in lakes the patterns are mainly related to thermal stratification and dissolved oxygen, nitrate and nitrite concentrations (Shade et al., 2007). Many of these variables are irrelevant to a subsurface system, and drivers of the variations were therefore different in our study. The oscillating dissimilarity patterns of the groundwater bacterial communities were largely explained by recharge that predominantly occurs during the winter half year (Lehmann & Totsche, 2020). Understanding the drivers and community reactions resulting in patterns unique to each aquatic system, will provide complementary insights into the wide range of processes shaping microbial communities across environments.

Microbial communities, in the absence of external influences, are most often shifting towards a stable composition; a change in this state can be brought about by changes in the conditions or a disturbance that pushes the system into a new state (Faust et al., 2015). Marine environments are a good example of a system where changes in the composition of microbial communities is driven by a change in the environmental conditions, and since these environmental conditions go through similar cycles each year, so do the communities. Seasonal patterns of marine bacterial communities are so consistent over the years that sampling month can be predicted based on bacterial community composition (Fuhrman et al., 2006). Lake bacterial communities are also largely shaped by gradually, and cyclically, changing environmental conditions, but they are more prone to disturbances. Heavy rainfall can result in mixing of the otherwise stratified water column in lakes, but lake microbial communities are resilient to such events, returning to their ‘pre-disturbance’ state once the environmental parameters also recover (Shade et al., 2012). Additionally, the bacterial community ‘recovery’ after rainfall related disturbances can be tracked through clear phases of succession and can occur as rapidly as within two weeks of the flooding event (Shabarova et al., 2021). These patterns are distinct to what we observed in our system, where the microbial communities never returned to the same state as observed during previously time points. This is likely because the temporal patterns in groundwater microbiomes are driven largely by disturbance, in the form of recharge events which impose ‘shocks’ on the aquifer communities. We found that these successive disturbances resulted in bacterial communities that were increasingly more dissimilar to their ‘initial’ state (i.e., at the first sampling point), and oscillating patterns were only apparent when comparing samples within a three-year time period. This suggests that both the type of disturbance as well as the system determines the microbial response.

In groundwater systems, the disturbances directly and repetitively introduce new surface derived taxa to a groundwater microbiome adapted to its local environmental conditions. We consider the shallow subsurface an open biogeoreactor, linked to the surface by water, matter and energy flow (Küsel et al., 2016) and the magnitude of the import of surface derived microbial taxa varies over space and time. Given its large biodiversity (Fierer et al., 2007), soils represent a rich microbial seedbank, and exported taxa will vary with each recharge event. This inconsistency will affect groundwater community assembly and might explain why unlike lake ecosystems groundwater microbiomes of this study did not return to the same state after each perturbation.

The Hainich CZE groundwater microbiome, dominated by Cand. Patescibacteria, Proteobacteria and Nitrospira, is distinct from the soil microbiome in the local recharge areas (Herrmann et al., 2019; Krüger et al., 2021). Acidobacteria, Actinobacteria, and Planctomycetes, making up approximately 40% of the soil bacterial community, are either not mobilized into the seepage at all, or in low abundance (Herrmann et al., 2019). Surprisingly, members of Cand. Patescibacteria already dominate seepage water at 30 cm soil depth with relative abundances of up to 50%, although they represent only 0.55% of the total soil community (Herrmann et al., 2019). Indeed, the Cand. Patescibacteria UBA9983, enriched in seepage compared to soil by a factor of 100 in that study, was among the recharge-favoured groundwater bacteria identified at well H41, in addition to other Patescibacteria such as Saccharimonadales and Ca. Peribacteria and members of the Proteobacteria phylum. Ultimately, the mobilized taxa become part of the present groundwater microbiome and either flourish, survive, starve, or die and end up as necromass for the survivors.

Downhill groundwater of well H52 is not characterized by high surface connectivity, and the contribution of recharge-related taxa was consistently small across the periods, and therefore not a dominant driver of the temporal patterns. This serves as a reminder that both dispersal and environmental selective community assembly mechanisms shape the dynamics of the three contrasting groundwater microbiomes present at the Hainich CZE. The groundwater microbiome at H52 is under stronger environmental selection pressure than those of the other two wells (Yan et al., 2020), due to longer groundwater residence time caused by lower bulk rock permeability than midslope wells (Lazar et al., 2019). As a result, the minor perturbation effects on the microbial community dynamics at H52 were further stabilized by higher selective assembly mechanisms leading to less dissimilar communities over time.

Surface export driven by recharge events alone did not explain the well-specific community dynamics we observed. Recharge events not only increase connectivity to surface soils but enable the exchange between different aquifers (Lehmann and Totsche, 2020), including the mobilization of rock-attached microbes and exchange with the groundwater. Surface attached communities have been shown to differ significantly to those found in the groundwater of the Hainich CZE (Lazar et al., 2017) and can therefore, like the soil, act as a microbial seedbank during recharge events. Larger recharge events would result in a greater exchange with these rock surface attached communities. Therefore, the large groundwater recharge observed in well H41 (> 20 m) could have resulted in greater dissemination of the microbial seedbank into the groundwater, resulting in the elevated alpha diversity we observed during recharge events. The large increase of alpha diversity following the pronounced water level increase in 2018 suggests a positive relationship between recharge strength and bacterial diversity in shallow aquifers with consequences for ecosystem functioning. As the provision of clean drinking water is one of the most important ecosystem services the subsurface provides to us humans, we have to study a wider range of groundwater systems to truly understand the temporal dynamics of groundwater microbiomes.

## Conclusions

The discrete but recurring disturbances, in the form or groundwater recharge, are an intrinsic system property of near-surface groundwater microbiomes. Importantly, the oscillating patterns in community similarity was only observed when comparing bacterial communities sampled within three years of each other. After this the temporal variation between samples was too great to observe the oscillating patterns, leading to decreased similarity to the initial state over time suggesting that community succession is occurring. The patterns we observed are unique to shallow groundwater environments and differ to what is seen in oceans, lakes and rivers aboveground, and highlight the importance of conducting high resolution temporal research across a wide variety of environments. Only then can we begin to achieve a complete understanding of the various factors and processes shaping microbial communities across the range of ecosystems they inhabit.

## Declarations

### Availability of data and material

The raw sequencing data were deposited in the European Sequence Archive (ENA), with BioProject IDs and accession numbers as shown in Supplementary Table S1. All R scripts for the statistical analyses are available at: https://github.com/pikatech/cyclic_patterns.

## Authors’ contributions

LY, KK and KT designed the research with input from MH and RL. LY performed the data analyses. LY and SH wrote the manuscript with input from all other co-authors.

## Acknowledgements

The authors wish to thank Heiko Minkmar, Bernd Ruppe, Falko Gutmann, Stefan Riedel and Patricia Geesink for groundwater sampling and on-site measurements/sample preparation. Special thanks are extended to Maria Fabisch for scientific coordination and Bernd Kampe for the help with AquaDiva literature research.

