## Supplementary for "Groundwater bacterial communities evolve over time, exhibiting oscillating similarity patterns in response to recharge"

**Supplementary Information**

Table S1. Summary of sequence data deposition in the European Nucleotide Archive (ENA)

| SampleID | Well | Samp_cam | Date_Ref | Type | Study accession no. | Sample accession no. | Reference |
|---|---|---|---|---|---|---|---|
| H41_PNK033 | H41 | PNK033 | 18/02/2013 | In-house | PRJEB35519 | ERS4043092 | Current study |
| H41_PNK034 | H41 | PNK034 | 18/03/2013 | In-house | PRJEB35519 | ERS4043093 | Current study |
| H41_PNK036 | H41 | PNK036 | 07/05/2013 | In-house | PRJEB35519 | ERS4043094 | Current study |
| H41_PNK038 | H41 | PNK038 | 03/07/2013 | In-house | PRJEB35519 | ERS4043095 | Current study |
| H41_PNK039 | H41 | PNK039 | 31/07/2013 | In-house | PRJEB35519 | ERS4043096 | Current study |
| H41_PNK040 | H41 | PNK040 | 28/08/2013 | In-house | PRJEB35519 | ERS4043097 | Current study |
| H41_PNK041 | H41 | PNK041 | 25/09/2013 | LGC genomics GmbH | PRJEB35519 | ERS4043098 | Current study |
| H41_PNK042 | H41 | PNK042 | 23/10/2013 | In-house | PRJEB35519 | ERS4043099 | Current study |
| H41_PNK043 | H41 | PNK043 | 20/11/2013 | In-house | PRJEB35519 | ERS4043100 | Current study |
| H41_PNK044 | H41 | PNK044 | 18/12/2013 | In-house | PRJEB35519 | ERS4043101 | Current study |
| H41_PNK045 | H41 | PNK045 | 15/01/2014 | In-house | PRJEB35519 | ERS4043102 | Current study |
| H41_PNK046 | H41 | PNK046 | 12/02/2014 | In-house | PRJEB35519 | ERS4043103 | Current study |
| H41_PNK047 | H41 | PNK047 | 12/03/2014 | In-house | PRJEB35519 | ERS4043104 | Current study |
| H41_PNK048 | H41 | PNK048 | 09/04/2014 | In-house | PRJEB35519 | ERS4043105 | Current study |
| H41_PNK050 | H41 | PNK050 | 04/06/2014 | In-house | PRJEB35519 | ERS4043106 | Current study |
| H41_PNK051 | H41 | PNK051 | 02/07/2014 | LGC genomics GmbH | PRJEB14968 | ERS1270620 | Schwab et al. (2017) |
| H41_PNK052 | H41 | PNK052 | 30/07/2014 | In-house | PRJEB35519 | ERS4043107 | Current study |
| H41_PNK053 | H41 | PNK053 | 27/08/2014 | LGC genomics GmbH | PRJEB35519 | ERS4043108 | Current study |
| H41_PNK054 | H41 | PNK054 | 24/09/2014 | LGC genomics GmbH | PRJEB33032 | ERS3518028 | Yan et al. (2020) |
| H41_PNK055 | H41 | PNK055 | 22/10/2014 | In-house | PRJEB35519 | ERS4043109 | Current study |
| H41_PNK056 | H41 | PNK056 | 19/11/2014 | In-house | PRJEB35519 | ERS4043110 | Current study |
| H41_PNK058 | H41 | PNK058 | 14/01/2015 | LGC genomics GmbH | PRJEB35519 | ERS4043111 | Current study |
| H41_PNK059 | H41 | PNK059 | 11/02/2015 | In-house | PRJEB35519 | ERS4043112 | Current study |
| H41_PNK060 | H41 | PNK060 | 10/03/2015 | LGC genomics GmbH | PRJEB33254 | ERS3548279 | Herrmann et al. (2019) |
| H41_PNK061 | H41 | PNK061 | 08/04/2015 | In-house | PRJEB35519 | ERS4043113 | Current study |
| H41_PNK062 | H41 | PNK062 | 05/05/2015 | In-house | PRJEB35519 | ERS4043114 | Current study |

| | | | | | | | |
|---|---|---|---|---|---|---|---|
| H41_PNK063 | H41 | PNK063 | 02/06/2015 | LGC genomics GmbH | PRJEB33254 | ERS3548281 | Herrmann et al. (2019) |
| H41_PNK064 | H41 | PNK064 | 29/06/2015 | In-house | PRJEB35519 | ERS4043115 | Current study |
| H41_PNK065 | H41 | PNK065 | 29/07/2015 | In-house | PRJEB35519 | ERS4043116 | Current study |
| H41_PNK066 | H41 | PNK066 | 25/08/2015 | LGC genomics GmbH | PRJEB20223 | ERS1645488 | Kumar et al. (2017) |
| H41_PNK067 | H41 | PNK067 | 23/09/2015 | LGC genomics GmbH | PRJEB25133 | ERS2221396 | Herrmann et al. (2019) |
| H41_PNK068 | H41 | PNK068 | 21/10/2015 | In-house | PRJEB35519 | ERS4043117 | Current study |
| H41_PNK069 | H41 | PNK069 | 18/11/2015 | LGC genomics GmbH | PRJEB20223 | ERS1645495 | Kumar et al. (2017) |
| H41_PNK070 | H41 | PNK070 | 15/12/2015 | LGC genomics GmbH | PRJEB25133 | ERS2221400 | Herrmann et al. (2019) |
| H41_PNK072 | H41 | PNK072 | 09/02/2016 | LGC genomics GmbH | PRJEB33032 | ERS3518029 | Yan et al. (2020) |
| H41_PNK073 | H41 | PNK073 | 08/03/2016 | In-house | PRJEB35519 | ERS4043118 | Current study |
| H41_PNK074 | H41 | PNK074 | 05/04/2016 | In-house | PRJEB35519 | ERS4043119 | Current study |
| H41_PNK075 | H41 | PNK075 | 03/05/2016 | In-house | PRJEB33032 | ERS3518030 | Yan et al. (2020) |
| H41_PNK076 | H41 | PNK076 | 01/06/2016 | LGC genomics GmbH | PRJEB25133 | ERS2221402 | Herrmann et al. (2019) |
| H41_PNK077 | H41 | PNK077 | 28/06/2016 | In-house | PRJEB35519 | ERS4043120 | Current study |
| H41_PNK078 | H41 | PNK078 | 26/07/2016 | LGC genomics GmbH | PRJEB25133 | ERS2221404 | Herrmann et al. (2019) |
| H41_PNK079 | H41 | PNK079 | 23/08/2016 | In-house | PRJEB33923 | ERS3646577 | Benk et al. (2019) |
| H41_PNK080 | H41 | PNK080 | 20/09/2016 | In-house | PRJEB35519 | ERS4043121 | Current study |
| H41_PNK081 | H41 | PNK081 | 18/10/2016 | LGC genomics GmbH | PRJEB25133 | ERS2221406 | Herrmann et al. (2019) |
| H41_PNK082 | H41 | PNK082 | 15/11/2016 | In-house | PRJEB33923 | ERS3646578 | Benk et al. (2019) |
| H41_PNK083 | H41 | PNK083 | 13/12/2016 | In-house | PRJEB35519 | ERS4043122 | Current study |
| H41_PNK084 | H41 | PNK084 | 10/01/2017 | LGC genomics GmbH | PRJEB33032 | ERS3518031 | Yan et al. (2020) |
| H41_PNK086 | H41 | PNK086 | 10/03/2017 | In-house | PRJEB33923 | ERS3646579 | Benk et al. (2019) |
| H41_PNK087 | H41 | PNK087 | 04/04/2017 | LGC genomics GmbH | PRJEB33032 | ERS3518032 | Yan et al. (2020) |
| H41_PNK088 | H41 | PNK088 | 05/05/2017 | In-house | PRJEB35519 | ERS4043123 | Current study |
| H41_PNK089 | H41 | PNK089 | 30/05/2017 | In-house | PRJEB33923 | ERS3646580 | Benk et al. (2019) |
| H41_PNK090 | H41 | PNK090 | 29/06/2017 | In-house | PRJEB35519 | ERS4043124 | Current study |
| H41_PNK091 | H41 | PNK091 | 28/07/2017 | In-house | PRJEB35519 | ERS4043125 | Current study |
| H41_PNK092 | H41 | PNK092 | 22/08/2017 | In-house | PRJEB35519 | ERS4043126 | Current study |
| H41_PNK094 | H41 | PNK094 | 17/10/2017 | In-house | PRJEB35519 | ERS4043127 | Current study |
| H41_PNK096 | H41 | PNK096 | 12/12/2017 | In-house | PRJEB35519 | ERS4043128 | Current study |
| H41_PNK097 | H41 | PNK097 | 09/01/2018 | In-house | PRJEB35519 | ERS4043129 | Current study |

| | | | | | | | |
|---|---|---|---|---|---|---|---|
| H41_PNK098 | H41 | PNK098 | 06/02/2018 | In-house | PRJEB35519 | ERS4043130 | Current study |
| H41_PNK099 | H41 | PNK099 | 07/03/2018 | In-house | PRJEB35519 | ERS4043131 | Current study |
| H41_PNK100 | H41 | PNK100 | 04/04/2018 | In-house | PRJEB35519 | ERS4043132 | Current study |
| H41_PNK101 | H41 | PNK101 | 09/05/2018 | In-house | PRJEB35519 | ERS4043133 | Current study |
| H41_PNK102 | H41 | PNK102 | 30/05/2018 | In-house | PRJEB35519 | ERS4043134 | Current study |
| H41_PNK103 | H41 | PNK103 | 27/06/2018 | In-house | PRJEB35519 | ERS4043135 | Current study |
| H41_PNK104 | H41 | PNK104 | 25/07/2018 | In-house | PRJEB35519 | ERS4043136 | Current study |
| H41_PNK105 | H41 | PNK105 | 22/08/2018 | In-house | PRJEB35519 | ERS4043137 | Current study |
| H41_PNK106 | H41 | PNK106 | 19/09/2018 | In-house | PRJEB35519 | ERS4043138 | Current study |
| H41_PNK107 | H41 | PNK107 | 17/10/2018 | In-house | PRJEB35519 | ERS4043139 | Current study |
| H41_PNK108 | H41 | PNK108 | 14/11/2018 | In-house | PRJEB35519 | ERS4043140 | Current study |
| H41_PNK109 | H41 | PNK109 | 12/12/2018 | In-house | PRJEB35519 | ERS4043141 | Current study |
| H41_PNK110 | H41 | PNK110 | 09/01/2019 | In-house | PRJEB35519 | ERS4043142 | Current study |
| H41_PNK111 | H41 | PNK111 | 06/02/2019 | In-house | PRJEB35519 | ERS4043143 | Current study |
| H41_PNK112 | H41 | PNK112 | 06/03/2019 | In-house | PRJEB35519 | ERS4043144 | Current study |
| H41_PNK113 | H41 | PNK113 | 03/04/2019 | In-house | PRJEB35519 | ERS4043145 | Current study |
| H41_PNK114 | H41 | PNK114 | 01/05/2019 | In-house | PRJEB35519 | ERS4043146 | Current study |
| H41_PNK115 | H41 | PNK115 | 29/05/2019 | In-house | PRJEB35519 | ERS4043147 | Current study |
| H43_PNK033 | H43 | PNK033 | 18/02/2013 | In-house | PRJEB35519 | ERS4043148 | Current study |
| H43_PNK034 | H43 | PNK034 | 18/03/2013 | In-house | PRJEB35519 | ERS4043149 | Current study |
| H43_PNK036 | H43 | PNK036 | 07/05/2013 | In-house | PRJEB35519 | ERS4043150 | Current study |
| H43_PNK038 | H43 | PNK038 | 03/07/2013 | In-house | PRJEB35519 | ERS4043151 | Current study |
| H43_PNK039 | H43 | PNK039 | 31/07/2013 | In-house | PRJEB35519 | ERS4043152 | Current study |
| H43_PNK040 | H43 | PNK040 | 28/08/2013 | In-house | PRJEB35519 | ERS4043153 | Current study |
| H43_PNK041 | H43 | PNK041 | 25/09/2013 | LGC genomics GmbH | PRJEB35519 | ERS4043154 | Current study |
| H43_PNK042 | H43 | PNK042 | 23/10/2013 | In-house | PRJEB35519 | ERS4043155 | Current study |
| H43_PNK043 | H43 | PNK043 | 20/11/2013 | In-house | PRJEB35519 | ERS4043156 | Current study |
| H43_PNK044 | H43 | PNK044 | 18/12/2013 | In-house | PRJEB35519 | ERS4043157 | Current study |
| H43_PNK045 | H43 | PNK045 | 15/01/2014 | In-house | PRJEB35519 | ERS4043158 | Current study |
| H43_PNK046 | H43 | PNK046 | 12/02/2014 | In-house | PRJEB35519 | ERS4043159 | Current study |
| H43_PNK047 | H43 | PNK047 | 12/03/2014 | In-house | PRJEB35519 | ERS4043160 | Current study |

| | | | | | | | |
|---|---|---|---|---|---|---|---|
| H43_PNK048 | H43 | PNK048 | 09/04/2014 | In-house | PRJEB35519 | ERS4043161 | Current study |
| H43_PNK049 | H43 | PNK049 | 08/05/2014 | In-house | PRJEB35519 | ERS4043162 | Current study |
| H43_PNK050 | H43 | PNK050 | 04/06/2014 | In-house | PRJEB35519 | ERS4043163 | Current study |
| H43_PNK051 | H43 | PNK051 | 02/07/2014 | LGC genomics GmbH | PRJEB14968 | ERS1270624 | Schwab et al. (2017) |
| H43_PNK052 | H43 | PNK052 | 30/07/2014 | In-house | PRJEB35519 | ERS4043164 | Current study |
| H43_PNK053 | H43 | PNK053 | 27/08/2014 | LGC genomics GmbH | PRJEB35519 | ERS4043165 | Current study |
| H43_PNK054 | H43 | PNK054 | 24/09/2014 | LGC genomics GmbH | PRJEB33032 | ERS3518038 | Yan et al. (2020) |
| H43_PNK055 | H43 | PNK055 | 22/10/2014 | In-house | PRJEB35519 | ERS4043166 | Current study |
| H43_PNK056 | H43 | PNK056 | 19/11/2014 | In-house | PRJEB35519 | ERS4043167 | Current study |
| H43_PNK058 | H43 | PNK058 | 14/01/2015 | LGC genomics GmbH | PRJEB35519 | ERS4043168 | Current study |
| H43_PNK059 | H43 | PNK059 | 11/02/2015 | In-house | PRJEB35519 | ERS4043169 | Current study |
| H43_PNK060 | H43 | PNK060 | 10/03/2015 | LGC genomics GmbH | PRJEB33254 | ERS3548289 | Herrmann et al. (2019) |
| H43_PNK061 | H43 | PNK061 | 08/04/2015 | In-house | PRJEB35519 | ERS4043170 | Current study |
| H43_PNK062 | H43 | PNK062 | 05/05/2015 | In-house | PRJEB35519 | ERS4043171 | Current study |
| H43_PNK063 | H43 | PNK063 | 02/06/2015 | LGC genomics GmbH | PRJEB33254 | ERS3548291 | Herrmann et al. (2019) |
| H43_PNK064 | H43 | PNK064 | 29/06/2015 | In-house | PRJEB35519 | ERS4043172 | Current study |
| H43_PNK065 | H43 | PNK065 | 29/07/2015 | In-house | PRJEB35519 | ERS4043173 | Current study |
| H43_PNK066 | H43 | PNK066 | 25/08/2015 | LGC genomics GmbH | PRJEB20223 | ERS1645490 | Kumar et al. (2017) |
| H43_PNK067 | H43 | PNK067 | 23/09/2015 | LGC genomics GmbH | PRJEB25133 | ERS2221420 | Herrmann et al. (2019) |
| H43_PNK068 | H43 | PNK068 | 21/10/2015 | In-house | PRJEB35519 | ERS4043174 | Current study |
| H43_PNK069 | H43 | PNK069 | 18/11/2015 | LGC genomics GmbH | PRJEB20223 | ERS1645497 | Kumar et al. (2017) |
| H43_PNK070 | H43 | PNK070 | 15/12/2015 | LGC genomics GmbH | PRJEB25133 | ERS2221424 | Herrmann et al. (2019) |
| H43_PNK072 | H43 | PNK072 | 09/02/2016 | LGC genomics GmbH | PRJEB33032 | ERS3518039 | Yan et al. (2020) |
| H43_PNK073 | H43 | PNK073 | 08/03/2016 | In-house | PRJEB35519 | ERS4043175 | Current study |
| H43_PNK074 | H43 | PNK074 | 05/04/2016 | In-house | PRJEB35519 | ERS4043176 | Current study |
| H43_PNK075 | H43 | PNK075 | 03/05/2016 | LGC genomics GmbH | PRJEB33032 | ERS3518040 | Yan et al. (2020) |
| H43_PNK076 | H43 | PNK076 | 01/06/2016 | LGC genomics GmbH | PRJEB25133 | ERS2221426 | Herrmann et al. (2019) |
| H43_PNK078 | H43 | PNK078 | 26/07/2016 | LGC genomics GmbH | PRJEB25133 | ERS2221428 | Herrmann et al. (2019) |
| H43_PNK079 | H43 | PNK079 | 23/08/2016 | In-house | PRJEB35519 | ERS4043177 | Current study |
| H43_PNK080 | H43 | PNK080 | 20/09/2016 | In-house | PRJEB35519 | ERS4043178 | Current study |
| H43_PNK081 | H43 | PNK081 | 18/10/2016 | LGC genomics GmbH | PRJEB25133 | ERS2221430 | Herrmann et al. (2019) |

| | | | | | | | |
|---|---|---|---|---|---|---|---|
| H43_PNK082 | H43 | PNK082 | 15/11/2016 | In-house | PRJEB35519 | ERS4043179 | Current study |
| H43_PNK083 | H43 | PNK083 | 13/12/2016 | In-house | PRJEB35519 | ERS4043180 | Current study |
| H43_PNK084 | H43 | PNK084 | 10/01/2017 | LGC genomics GmbH | PRJEB33032 | ERS3518041 | Yan et al. (2020) |
| H43_PNK086 | H43 | PNK086 | 10/03/2017 | In-house | PRJEB35519 | ERS4043181 | Current study |
| H43_PNK087 | H43 | PNK087 | 04/04/2017 | LGC genomics GmbH | PRJEB33032 | ERS3518042 | Yan et al. (2020) |
| H43_PNK088 | H43 | PNK088 | 05/05/2017 | In-house | PRJEB35519 | ERS4043182 | Current study |
| H43_PNK089 | H43 | PNK089 | 30/05/2017 | In-house | PRJEB35519 | ERS4043183 | Current study |
| H43_PNK090 | H43 | PNK090 | 29/06/2017 | In-house | PRJEB35519 | ERS4043184 | Current study |
| H43_PNK091 | H43 | PNK091 | 28/07/2017 | In-house | PRJEB35519 | ERS4043185 | Current study |
| H43_PNK092 | H43 | PNK092 | 22/08/2017 | In-house | PRJEB35519 | ERS4043186 | Current study |
| H43_PNK094 | H43 | PNK094 | 17/10/2017 | In-house | PRJEB35519 | ERS4043187 | Current study |
| H43_PNK096 | H43 | PNK096 | 12/12/2017 | In-house | PRJEB35519 | ERS4043188 | Current study |
| H43_PNK097 | H43 | PNK097 | 09/01/2018 | In-house | PRJEB35519 | ERS4043189 | Current study |
| H43_PNK098 | H43 | PNK098 | 06/02/2018 | In-house | PRJEB35519 | ERS4043190 | Current study |
| H43_PNK099 | H43 | PNK099 | 07/03/2018 | In-house | PRJEB35519 | ERS4043191 | Current study |
| H43_PNK100 | H43 | PNK100 | 04/04/2018 | In-house | PRJEB35519 | ERS4043192 | Current study |
| H43_PNK101 | H43 | PNK101 | 09/05/2018 | In-house | PRJEB35519 | ERS4043193 | Current study |
| H43_PNK102 | H43 | PNK102 | 30/05/2018 | In-house | PRJEB35519 | ERS4043194 | Current study |
| H43_PNK103 | H43 | PNK103 | 27/06/2018 | In-house | PRJEB35519 | ERS4043195 | Current study |
| H43_PNK104 | H43 | PNK104 | 25/07/2018 | In-house | PRJEB35519 | ERS4043196 | Current study |
| H43_PNK105 | H43 | PNK105 | 22/08/2018 | In-house | PRJEB35519 | ERS4043197 | Current study |
| H43_PNK106 | H43 | PNK106 | 19/09/2018 | In-house | PRJEB35519 | ERS4043198 | Current study |
| H43_PNK107 | H43 | PNK107 | 17/10/2018 | In-house | PRJEB35519 | ERS4043199 | Current study |
| H43_PNK108 | H43 | PNK108 | 14/11/2018 | In-house | PRJEB35519 | ERS4043200 | Current study |
| H43_PNK109 | H43 | PNK109 | 12/12/2018 | In-house | PRJEB35519 | ERS4043201 | Current study |
| H43_PNK110 | H43 | PNK110 | 09/01/2019 | In-house | PRJEB35519 | ERS4043202 | Current study |
| H43_PNK111 | H43 | PNK111 | 06/02/2019 | In-house | PRJEB35519 | ERS4043203 | Current study |
| H43_PNK112 | H43 | PNK112 | 06/03/2019 | In-house | PRJEB35519 | ERS4043204 | Current study |
| H43_PNK113 | H43 | PNK113 | 03/04/2019 | In-house | PRJEB35519 | ERS4043205 | Current study |
| H43_PNK114 | H43 | PNK114 | 01/05/2019 | In-house | PRJEB35519 | ERS4043206 | Current study |
| H43_PNK115 | H43 | PNK115 | 29/05/2019 | In-house | PRJEB35519 | ERS4043207 | Current study |

| | | | | | | | |
|---|---|---|---|---|---|---|---|
| H52_PNK036 | H52 | PNK036 | 07/05/2013 | In-house | PRJEB35519 | ERS4043208 | Current study |
| H52_PNK037 | H52 | PNK037 | 05/06/2013 | In-house | PRJEB35519 | ERS4043209 | Current study |
| H52_PNK038 | H52 | PNK038 | 03/07/2013 | In-house | PRJEB35519 | ERS4043210 | Current study |
| H52_PNK039 | H52 | PNK039 | 31/07/2013 | In-house | PRJEB35519 | ERS4043211 | Current study |
| H52_PNK040 | H52 | PNK040 | 28/08/2013 | In-house | PRJEB35519 | ERS4043212 | Current study |
| H52_PNK041 | H52 | PNK041 | 25/09/2013 | LGC genomics GmbH | PRJEB35519 | ERS4043213 | Current study |
| H52_PNK042 | H52 | PNK042 | 23/10/2013 | In-house | PRJEB35519 | ERS4043214 | Current study |
| H52_PNK043 | H52 | PNK043 | 20/11/2013 | In-house | PRJEB35519 | ERS4043215 | Current study |
| H52_PNK044 | H52 | PNK044 | 18/12/2013 | In-house | PRJEB35519 | ERS4043216 | Current study |
| H52_PNK045 | H52 | PNK045 | 15/01/2014 | In-house | PRJEB35519 | ERS4043217 | Current study |
| H52_PNK046 | H52 | PNK046 | 12/02/2014 | In-house | PRJEB35519 | ERS4043218 | Current study |
| H52_PNK047 | H52 | PNK047 | 12/03/2014 | In-house | PRJEB35519 | ERS4043219 | Current study |
| H52_PNK048 | H52 | PNK048 | 09/04/2014 | In-house | PRJEB35519 | ERS4043220 | Current study |
| H52_PNK049 | H52 | PNK049 | 08/05/2014 | In-house | PRJEB35519 | ERS4043221 | Current study |
| H52_PNK050 | H52 | PNK050 | 04/06/2014 | In-house | PRJEB35519 | ERS4043222 | Current study |
| H52_PNK051 | H52 | PNK051 | 02/07/2014 | LGC genomics GmbH | PRJEB14968 | ERS1270628 | Schwab et al. (2017) |
| H52_PNK052 | H52 | PNK052 | 30/07/2014 | In-house | PRJEB35519 | ERS4043223 | Current study |
| H52_PNK053 | H52 | PNK053 | 27/08/2014 | LGC genomics GmbH | PRJEB35519 | ERS4043224 | Current study |
| H52_PNK054 | H52 | PNK054 | 24/09/2014 | LGC genomics GmbH | PRJEB33032 | ERS3518046 | Yan et al. (2020) |
| H52_PNK055 | H52 | PNK055 | 22/10/2014 | In-house | PRJEB35519 | ERS4043225 | Current study |
| H52_PNK056 | H52 | PNK056 | 19/11/2014 | In-house | PRJEB35519 | ERS4043226 | Current study |
| H52_PNK057 | H52 | PNK057 | 16/12/2014 | LGC genomics GmbH | PRJEB33254 | ERS3548300 | Herrmann et al. (2019) |
| H52_PNK058 | H52 | PNK058 | 14/01/2015 | LGC genomics GmbH | PRJEB35519 | ERS4043227 | Current study |
| H52_PNK059 | H52 | PNK059 | 11/02/2015 | In-house | PRJEB35519 | ERS4043228 | Current study |
| H52_PNK060 | H52 | PNK060 | 10/03/2015 | LGC genomics GmbH | PRJEB33254 | ERS3548301 | Herrmann et al. (2019) |
| H52_PNK061 | H52 | PNK061 | 08/04/2015 | In-house | PRJEB35519 | ERS4043229 | Current study |
| H52_PNK062 | H52 | PNK062 | 05/05/2015 | In-house | PRJEB35519 | ERS4043230 | Current study |
| H52_PNK063 | H52 | PNK063 | 02/06/2015 | LGC genomics GmbH | PRJEB33254 | ERS3548303 | Herrmann et al. (2019) |
| H52_PNK064 | H52 | PNK064 | 29/06/2015 | In-house | PRJEB35519 | ERS4043231 | Current study |
| H52_PNK065 | H52 | PNK065 | 29/07/2015 | In-house | PRJEB35519 | ERS4043232 | Current study |
| H52_PNK066 | H52 | PNK066 | 25/08/2015 | LGC genomics GmbH | PRJEB20223 | ERS1645492 | Kumar et al. (2017) |

| | | | | | | | |
|---|---|---|---|---|---|---|---|
| H52_PNK067 | H52 | PNK067 | 23/09/2015 | LGC genomics GmbH | PRJEB25133 | ERS2221444 | Herrmann et al. (2019) |
| H52_PNK068 | H52 | PNK068 | 21/10/2015 | In-house | PRJEB35519 | ERS4043233 | Current study |
| H52_PNK069 | H52 | PNK069 | 18/11/2015 | LGC genomics GmbH | PRJEB20223 | ERS1645499 | Kumar et al. (2017) |
| H52_PNK070 | H52 | PNK070 | 15/12/2015 | LGC genomics GmbH | PRJEB25133 | ERS2221448 | Herrmann et al. (2019) |
| H52_PNK071 | H52 | PNK071 | 12/01/2016 | In-house | PRJEB35519 | ERS4043234 | Current study |
| H52_PNK072 | H52 | PNK072 | 09/02/2016 | LGC genomics GmbH | PRJEB33032 | ERS3518047 | Yan et al. (2020) |
| H52_PNK073 | H52 | PNK073 | 08/03/2016 | In-house | PRJEB35519 | ERS4043235 | Current study |
| H52_PNK074 | H52 | PNK074 | 05/04/2016 | In-house | PRJEB35519 | ERS4043236 | Current study |
| H52_PNK075 | H52 | PNK075 | 03/05/2016 | LGC genomics GmbH | PRJEB26565 | ERS2461671 | Lazar et al. (2019) |
| H52_PNK076 | H52 | PNK076 | 01/06/2016 | LGC genomics GmbH | PRJEB25133 | ERS2221450 | Herrmann et al. (2019) |
| H52_PNK077 | H52 | PNK077 | 28/06/2016 | In-house | PRJEB35519 | ERS4043237 | Current study |
| H52_PNK078 | H52 | PNK078 | 26/07/2016 | LGC genomics GmbH | PRJEB25133 | ERS2221452 | Herrmann et al. (2019) |
| H52_PNK079 | H52 | PNK079 | 23/08/2016 | In-house | PRJEB35519 | ERS4043238 | Current study |
| H52_PNK080 | H52 | PNK080 | 20/09/2016 | In-house | PRJEB35519 | ERS4043239 | Current study |
| H52_PNK081 | H52 | PNK081 | 18/10/2016 | LGC genomics GmbH | PRJEB25133 | ERS2221454 | Herrmann et al. (2019) |
| H52_PNK082 | H52 | PNK082 | 15/11/2016 | In-house | PRJEB35519 | ERS4043240 | Current study |
| H52_PNK083 | H52 | PNK083 | 13/12/2016 | In-house | PRJEB35519 | ERS4043241 | Current study |
| H52_PNK084 | H52 | PNK084 | 10/01/2017 | LGC genomics GmbH | PRJEB33032 | ERS3518048 | Yan et al. (2020) |
| H52_PNK085 | H52 | PNK085 | 04/02/2017 | In-house | PRJEB35519 | ERS4043242 | Current study |
| H52_PNK086 | H52 | PNK086 | 10/03/2017 | In-house | PRJEB35519 | ERS4043243 | Current study |
| H52_PNK087 | H52 | PNK087 | 04/04/2017 | LGC genomics GmbH | PRJEB33032 | ERS3518049 | Yan et al. (2020) |
| H52_PNK088 | H52 | PNK088 | 05/05/2017 | In-house | PRJEB35519 | ERS4043244 | Current study |
| H52_PNK089 | H52 | PNK089 | 30/05/2017 | In-house | PRJEB35519 | ERS4043245 | Current study |
| H52_PNK090 | H52 | PNK090 | 29/06/2017 | In-house | PRJEB35519 | ERS4043246 | Current study |
| H52_PNK091 | H52 | PNK091 | 28/07/2017 | In-house | PRJEB35519 | ERS4043247 | Current study |
| H52_PNK092 | H52 | PNK092 | 22/08/2017 | In-house | PRJEB35519 | ERS4043248 | Current study |
| H52_PNK093 | H52 | PNK093 | 19/09/2017 | In-house | PRJEB35519 | ERS4043249 | Current study |
| H52_PNK094 | H52 | PNK094 | 17/10/2017 | In-house | PRJEB35519 | ERS4043250 | Current study |
| H52_PNK095 | H52 | PNK095 | 13/11/2017 | In-house | PRJEB35519 | ERS4043251 | Current study |
| H52_PNK096 | H52 | PNK096 | 12/12/2017 | In-house | PRJEB35519 | ERS4043252 | Current study |
| H52_PNK097 | H52 | PNK097 | 09/01/2018 | In-house | PRJEB35519 | ERS4043253 | Current study |

| | | | | | | | |
|---|---|---|---|---|---|---|---|
| H52_PNK098 | H52 | PNK098 | 06/02/2018 | In-house | PRJEB35519 | ERS4043254 | Current study |
| H52_PNK099 | H52 | PNK099 | 07/03/2018 | In-house | PRJEB35519 | ERS4043255 | Current study |
| H52_PNK100 | H52 | PNK100 | 04/04/2018 | In-house | PRJEB35519 | ERS4043257 | Current study |
| H52_PNK101 | H52 | PNK101 | 09/05/2018 | In-house | PRJEB35519 | ERS4043258 | Current study |
| H52_PNK102 | H52 | PNK102 | 30/05/2018 | In-house | PRJEB35519 | ERS4043259 | Current study |
| H52_PNK103 | H52 | PNK103 | 27/06/2018 | In-house | PRJEB35519 | ERS4043260 | Current study |
| H52_PNK104 | H52 | PNK104 | 25/07/2018 | In-house | PRJEB35519 | ERS4043261 | Current study |
| H52_PNK105 | H52 | PNK105 | 22/08/2018 | In-house | PRJEB35519 | ERS4043262 | Current study |
| H52_PNK106 | H52 | PNK106 | 19/09/2018 | In-house | PRJEB35519 | ERS4043263 | Current study |
| H52_PNK107 | H52 | PNK107 | 17/10/2018 | In-house | PRJEB35519 | ERS4043264 | Current study |
| H52_PNK108 | H52 | PNK108 | 14/11/2018 | In-house | PRJEB35519 | ERS4043265 | Current study |
| H52_PNK109 | H52 | PNK109 | 12/12/2018 | In-house | PRJEB35519 | ERS4043266 | Current study |
| H52_PNK110 | H52 | PNK110 | 09/01/2019 | In-house | PRJEB35519 | ERS4043267 | Current study |
| H52_PNK111 | H52 | PNK111 | 06/02/2019 | In-house | PRJEB35519 | ERS4043268 | Current study |
| H52_PNK112 | H52 | PNK112 | 06/03/2019 | In-house | PRJEB35519 | ERS4043269 | Current study |
| H52_PNK113 | H52 | PNK113 | 03/04/2019 | In-house | PRJEB35519 | ERS4043270 | Current study |
| H52_PNK114 | H52 | PNK114 | 01/05/2019 | In-house | PRJEB35519 | ERS4043271 | Current study |
| H52_PNK115 | H52 | PNK115 | 29/05/2019 | In-house | PRJEB35519 | ERS4043272 | Current study |

Table S2. Relative abundance (%) of each bacterial class and phylum in the groundwater of H41.

| Phylum | Class | Mean (%) | Standard deviation (%) |
|---|---|---|---|
| Acetothermia | Acetothermiia | 0.004 | 0.006 |
| Acidobacteria | | 2.068 | 0.874 |
| | Subgroup_6 | 0.505 | 0.659 |
| | Blastocatellia_(Subgroup_4) | 0.408 | 0.354 |
| | Acidobacteriia | 0.312 | 0.178 |
| | Holophagae | 0.257 | 0.133 |
| | Thermoanaerobaculia | 0.240 | 0.133 |
| | Subgroup_22 | 0.171 | 0.086 |
| | Subgroup_17 | 0.117 | 0.101 |
| | Subgroup_15 | 0.014 | 0.017 |
| | AT-s3-28 | 0.013 | 0.017 |
| | Subgroup_5 | 0.008 | 0.014 |
| | Subgroup_20 | 0.005 | 0.008 |
| | Acidobacteria_unclassified | 0.004 | 0.006 |
| | Subgroup_21 | 0.004 | 0.009 |
| | Subgroup_11 | 0.004 | 0.008 |
| | Subgroup_19 | 0.003 | 0.006 |
| | Aminicenantia | 0.002 | 0.005 |
| | Subgroup_18 | 0.001 | 0.007 |
| | Subgroup_9 | 0.001 | 0.003 |
| Actinobacteria | | 2.367 | 1.378 |
| | Actinobacteria | 0.855 | 0.792 |
| | Acidimicrobiia | 0.846 | 0.499 |
| | Thermoleophilia | 0.491 | 0.345 |
| | MB-A2-108 | 0.063 | 0.061 |
| | Coriobacteriia | 0.053 | 0.044 |
| | Actinobacteria_unclassified | 0.048 | 0.067 |
| | WCHB1-81 | 0.006 | 0.011 |
| | RBG-16-55-12 | 0.004 | 0.019 |
| Armatimonadetes | | 0.073 | 0.056 |
| | Fimbriimonadia | 0.071 | 0.056 |
| | uncultured | 0.002 | 0.004 |
| | Chthonomonadetes | 0.001 | 0.002 |
| Bacteria_unclassified | Bacteria_unclassified | 4.912 | 2.001 |
| Bacteroidetes | | 3.198 | 1.970 |
| | Bacteroidia | 2.893 | 1.961 |
| | Ignavibacteria | 0.301 | 0.139 |
| | OC31 | 0.004 | 0.016 |
| BHI80-139 | BHI80-139_cl | 0.005 | 0.012 |
| BRC1 | BRC1_cl | 0.022 | 0.033 |
| Calditrichaeota | Calditrichia | 0.002 | 0.008 |
| Chlamydiae | Chlamydiae | 0.266 | 0.431 |
| Chloroflexi | | 3.700 | 1.646 |
| | KD4-96 | 1.138 | 0.715 |

| | | | |
|---|---|---|---|
| | Dehalococcoidia | 1.112 | 0.556 |
| | Anaerolineae | 0.522 | 0.351 |
| | JG30-KF-CM66 | 0.484 | 0.325 |
| | Chloroflexi_unclassified | 0.207 | 0.141 |
| | Gitt-GS-136 | 0.080 | 0.054 |
| | OLB14 | 0.068 | 0.067 |
| | TK10 | 0.034 | 0.041 |
| | P2-11E | 0.031 | 0.028 |
| | Chloroflexia | 0.023 | 0.058 |
| | TK17 | 0.002 | 0.005 |
| Cyanobacteria | | 0.287 | 0.193 |
| | Sericytochromatia | 0.202 | 0.186 |
| | Melainabacteria | 0.063 | 0.099 |
| | Cyanobacteria_unclassified | 0.020 | 0.029 |
| | Oxyphotobacteria | 0.001 | 0.005 |
| Dadabacteria | Dadabacteriia | 0.008 | 0.010 |
| Deinococcus-Thermus | Deinococci | 0.007 | 0.023 |
| Dependentiae | Babeliae | 0.234 | 0.120 |
| Elusimicrobia | | 1.140 | 0.596 |
| | Elusimicrobia | 1.097 | 0.593 |
| | Lineage_IIc | 0.023 | 0.031 |
| | Endomicrobia | 0.007 | 0.040 |
| | Lineage_IIb | 0.005 | 0.009 |
| | Lineage_IIa | 0.005 | 0.010 |
| | Elusimicrobia_cl | 0.003 | 0.005 |
| Entotheonellaeota | Entotheonellia | 0.014 | 0.019 |
| Epsilonbacteraeota | Campylobacteria | 0.196 | 0.677 |
| FCPU426 | FCPU426_cl | 0.021 | 0.035 |
| Fibrobacteres | | 0.123 | 0.194 |
| | Fibrobacteria | 0.105 | 0.197 |
| | Chitinivibrionia | 0.018 | 0.021 |
| Firestonebacteria | Firestonebacteria_cl | 0.004 | 0.008 |
| Firmicutes | | 0.259 | 0.363 |
| | Clostridia | 0.215 | 0.296 |
| | Bacilli | 0.037 | 0.121 |
| | Negativicutes | 0.007 | 0.015 |
| Fusobacteria | Fusobacteriia | 0.009 | 0.021 |
| GAL15 | GAL15_cl | 0.106 | 0.083 |
| Gemmatimonadetes | | 1.498 | 0.864 |
| | Gemmatimonadetes | 1.487 | 0.862 |
| | BD2-11_terrestrial_group | 0.005 | 0.010 |
| | Longimicrobia | 0.004 | 0.009 |
| | MD2902-B12 | 0.002 | 0.004 |
| Hydrogenedentes | Hydrogenedentia | 0.097 | 0.055 |
| Kiritimatiellaeota | Kiritimatiellae | 0.030 | 0.084 |
| Latescibacteria | | 0.814 | 0.371 |

| | | | |
|---|---|---|---|
| | Latescibacteria_cl | 0.805 | 0.365 |
| | Latescibacteria | 0.010 | 0.016 |
| Lentisphaerae | | 0.068 | 0.097 |
| | Lentisphaeria | 0.057 | 0.071 |
| | Oligosphaeria | 0.007 | 0.028 |
| | Lentisphaerae_unclassified | 0.004 | 0.013 |
| Lindowbacteria | Lindowbacteria_cl | 0.016 | 0.017 |
| Margulisbacteria | Margulisbacteria_cl | 0.076 | 0.039 |
| Nitrospinae | | 0.240 | 0.213 |
| | Nitrospinia | 0.209 | 0.198 |
| | P9X2b3D02 | 0.027 | 0.062 |
| | MT5B39 | 0.004 | 0.013 |
| Nitrospirae | | 14.083 | 8.422 |
| | Nitrospira | 9.422 | 5.736 |
| | Thermodesulfovibrionia | 4.563 | 3.263 |
| | 4-29-1 | 0.084 | 0.125 |
| | HDB-SIOI1093 | 0.009 | 0.012 |
| | Nitrospirae_unclassified | 0.005 | 0.010 |
| Omnitrophicaeota | | 3.116 | 2.192 |
| | Omnitrophicaeota_cl | 2.751 | 1.901 |
| | Omnitrophia | 0.357 | 0.325 |
| | Omnitrophicaeota_unclassified | 0.008 | 0.010 |
| Patescibacteria | | 18.315 | 9.156 |
| | Parcubacteria | 11.603 | 7.745 |
| | ABY1 | 3.295 | 1.432 |
| | Gracilibacteria | 1.732 | 0.884 |
| | Saccharimonadia | 1.268 | 0.961 |
| | Patescibacteria_unclassified | 0.239 | 0.274 |
| | Microgenomatia | 0.069 | 0.401 |
| | CPR2 | 0.055 | 0.097 |
| | Berkelbacteria | 0.054 | 0.065 |
| | Kazania | 0.001 | 0.005 |
| PAUC34f | PAUC34f_cl | 0.005 | 0.008 |
| Planctomycetes | | 6.009 | 2.427 |
| | Phycisphaerae | 1.905 | 1.166 |
| | Brocadiae | 1.729 | 1.197 |
| | OM190 | 0.972 | 0.633 |
| | Planctomycetacia | 0.550 | 0.523 |
| | vadinHA49 | 0.361 | 0.236 |
| | Pla4_lineage | 0.243 | 0.177 |
| | Planctomycetes_unclassified | 0.103 | 0.090 |
| | BD7-11 | 0.098 | 0.083 |
| | Pla3_lineage | 0.041 | 0.034 |
| | 028H05-P-BN-P5 | 0.007 | 0.009 |
| Poribacteria | Poribacteria_cl | 0.003 | 0.006 |
| Proteobacteria | | 34.771 | 10.148 |

| | | | |
|---|---|---|---|
| | Gammaproteobacteria | 16.635 | 6.921 |
| | Deltaproteobacteria | 9.344 | 3.664 |
| | Alphaproteobacteria | 8.361 | 6.398 |
| | Proteobacteria_unclassified | 0.429 | 0.241 |
| | Magnetococcia | 0.002 | 0.006 |
| Rokubacteria | NC10 | 0.637 | 0.370 |
| Schekmanbacteria | Schekmanbacteria_cl | 0.002 | 0.006 |
| Spirochaetes | | 0.160 | 0.226 |
| | MVP-15 | 0.109 | 0.226 |
| | Leptospirae | 0.029 | 0.029 |
| | Spirochaetia | 0.019 | 0.025 |
| | uncultured | 0.003 | 0.011 |
| Tenericutes | Mollicutes | 0.051 | 0.296 |
| Verrucomicrobia | Verrucomicrobiae | 0.526 | 0.322 |
| WOR-1 | WOR-1_cl | 0.034 | 0.035 |
| WPS-2 | WPS-2_cl | 0.129 | 0.121 |
| Zixibacteria | Zixibacteria_cl | 0.296 | 0.141 |

Table S3. Relative abundance (%) of each bacterial class and phylum in the groundwater of H43.

| Phylum | Class | Mean (%) | Standard deviation (%) |
|---|---|---|---|
| Acidobacteria | | 3,092 | 2,683 |
| | Subgroup_6 | 1,721 | 2,412 |
| | Acidobacteriia | 0,831 | 0,459 |
| | Holophagae | 0,149 | 0,142 |
| | Aminicenantia | 0,104 | 0,067 |
| | Blastocatellia_(Subgroup_4) | 0,068 | 0,048 |
| | Thermoanaerobaculia | 0,048 | 0,078 |
| | Subgroup_22 | 0,039 | 0,035 |
| | Subgroup_15 | 0,039 | 0,039 |
| | Subgroup_17 | 0,032 | 0,034 |
| | Subgroup_19 | 0,020 | 0,031 |
| | Subgroup_18 | 0,019 | 0,023 |
| | Subgroup_20 | 0,010 | 0,013 |
| | Subgroup_5 | 0,007 | 0,011 |
| | Subgroup_25 | 0,002 | 0,004 |
| | Acidobacteria_unclassified | 0,002 | 0,004 |
| Actinobacteria | | 2,433 | 1,925 |
| | Actinobacteria | 1,355 | 1,053 |
| | Thermoleophilia | 0,517 | 0,940 |
| | Acidimicrobiia | 0,365 | 0,197 |
| | MB-A2-108 | 0,093 | 0,087 |
| | RBG-16-55-12 | 0,036 | 0,082 |
| | WCHB1-81 | 0,029 | 0,031 |
| | Actinobacteria_unclassified | 0,022 | 0,039 |
| | Coriobacteriia | 0,016 | 0,015 |
| AncK6 | AncK6_cl | 0,012 | 0,015 |
| Armatimonadetes | | 0,054 | 0,041 |
| | Fimbriimonadia | 0,041 | 0,033 |
| | DG-56 | 0,006 | 0,012 |
| | Chthonomonadetes | 0,004 | 0,011 |
| | uncultured | 0,002 | 0,007 |
| Atribacteria | JS1 | 0,001 | 0,004 |
| Bacteria_unclassified | Bacteria_unclassified | 5,568 | 1,962 |
| Bacteroidetes | | 3,210 | 1,537 |
| | Bacteroidia | 2,550 | 1,250 |
| | Ignavibacteria | 0,653 | 0,618 |
| | Rhodothermia | 0,007 | 0,008 |
| BRC1 | BRC1_cl | 0,040 | 0,037 |
| Calditrichaeota | Calditrichia | 0,002 | 0,003 |
| Chlamydiae | | 1,686 | 1,470 |
| | Chlamydiae | 1,680 | 1,473 |
| | LD1-PA32 | 0,007 | 0,015 |
| Chloroflexi | | 5,000 | 2,224 |
| | Dehalococcoidia | 2,817 | 1,662 |

| | | | |
|---|---|---|---|
| | Anaerolineae | 0,884 | 0,459 |
| | Chloroflexi_unclassified | 0,527 | 0,305 |
| | KD4-96 | 0,298 | 0,211 |
| | JG30-KF-CM66 | 0,210 | 0,105 |
| | OLB14 | 0,137 | 0,138 |
| | P2-11E | 0,083 | 0,082 |
| | Chloroflexia | 0,027 | 0,050 |
| | Gitt-GS-136 | 0,012 | 0,021 |
| | TK10 | 0,003 | 0,010 |
| | AD3 | 0,002 | 0,005 |
| CK-2C2-2 | CK-2C2-2_cl | 0,005 | 0,010 |
| Cyanobacteria | | 0,189 | 0,137 |
| | Melainabacteria | 0,112 | 0,092 |
| | Cyanobacteria_unclassified | 0,040 | 0,064 |
| | Sericytochromatia | 0,036 | 0,046 |
| Dadabacteria | Dadabacteriia | 0,001 | 0,003 |
| Deinococcus-Thermus | Deinococci | 0,004 | 0,013 |
| Dependentiae | Babeliae | 0,576 | 0,309 |
| Edwardsbacteria | Edwardsbacteria_cl | 0,001 | 0,003 |
| Elusimicrobia | | 1,061 | 0,565 |
| | Elusimicrobia | 0,941 | 0,510 |
| | Lineage_IIc | 0,040 | 0,061 |
| | Elusimicrobia_cl | 0,035 | 0,037 |
| | Lineage_IIb | 0,022 | 0,022 |
| | Lineage_IIa | 0,013 | 0,020 |
| | Endomicrobia | 0,009 | 0,023 |
| | Elusimicrobia_unclassified | 0,001 | 0,002 |
| Epsilonbacteraeota | Campylobacteria | 0,171 | 0,501 |
| FCPU426 | FCPU426_cl | 0,012 | 0,016 |
| Fibrobacteres | | 0,117 | 0,098 |
| | Fibrobacteria | 0,102 | 0,094 |
| | Chitinivibrionia | 0,014 | 0,023 |
| Firestonebacteria | Firestonebacteria_cl | 0,005 | 0,007 |
| Firmicutes | | 1,004 | 1,069 |
| | Clostridia | 0,892 | 1,030 |
| | Bacilli | 0,076 | 0,072 |
| | Firmicutes_unclassified | 0,024 | 0,029 |
| | Negativicutes | 0,009 | 0,050 |
| | Erysipelotrichia | 0,003 | 0,009 |
| Fusobacteria | Fusobacteriia | 0,033 | 0,059 |
| GAL15 | GAL15_cl | 0,099 | 0,078 |
| Gemmatimonadetes | | 0,811 | 0,459 |
| | Gemmatimonadetes | 0,725 | 0,436 |
| | BD2-11_terrestrial_group | 0,062 | 0,044 |
| | MD2902-B12 | 0,018 | 0,020 |
| | Longimicrobia | 0,005 | 0,015 |

| | | | |
|---|---|---|---|
| Hydrogenedentes | Hydrogenedentia | 0,064 | 0,052 |
| Kiritimatiellaeota | Kiritimatiellae | 1,165 | 0,633 |
| Latescibacteria | | 0,511 | 0,293 |
| | Latescibacteria_cl | 0,488 | 0,282 |
| | Latescibacteria | 0,023 | 0,021 |
| LCP-89 | LCP-89_cl | 0,002 | 0,005 |
| Lentisphaerae | | 0,306 | 0,236 |
| | Lentisphaeria | 0,165 | 0,112 |
| | Oligosphaeria | 0,127 | 0,165 |
| | Lentisphaerae_unclassified | 0,014 | 0,022 |
| Lindowbacteria | Lindowbacteria_cl | 0,002 | 0,004 |
| Margulisbacteria | Margulisbacteria_cl | 0,090 | 0,062 |
| Nitrospinae | | 0,196 | 0,406 |
| | P9X2b3D02 | 0,111 | 0,418 |
| | Nitrospinia | 0,072 | 0,050 |
| | MT5B39 | 0,013 | 0,014 |
| Nitrospirae | | 2,347 | 2,091 |
| | Thermodesulfovibrionia | 1,473 | 1,707 |
| | Nitrospira | 0,796 | 1,028 |
| | 4-29-1 | 0,057 | 0,058 |
| | Nitrospirae_unclassified | 0,021 | 0,026 |
| Omnitrophicaeota | | 4,130 | 2,522 |
| | Omnitrophicaeota_cl | 3,803 | 2,358 |
| | Omnitrophia | 0,327 | 0,242 |
| Patescibacteria | | 26,348 | 12,693 |
| | Parcubacteria | 16,856 | 10,082 |
| | ABY1 | 5,104 | 2,449 |
| | Saccharimonadia | 1,638 | 1,738 |
| | Gracilibacteria | 1,210 | 0,531 |
| | Berkelbacteria | 1,090 | 0,863 |
| | Patescibacteria_unclassified | 0,256 | 0,156 |
| | CPR2 | 0,148 | 0,362 |
| | Microgenomatia | 0,033 | 0,069 |
| | Kazania | 0,013 | 0,018 |
| | uncultured | 0,002 | 0,004 |
| PAUC34f | PAUC34f_cl | 0,010 | 0,011 |
| Planctomycetes | | 3,301 | 1,729 |
| | Brocadiae | 1,165 | 0,871 |
| | Phycisphaerae | 1,164 | 1,172 |
| | Planctomycetacia | 0,264 | 0,200 |
| | vadinHA49 | 0,219 | 0,165 |
| | OM190 | 0,208 | 0,153 |
| | Pla4_lineage | 0,128 | 0,107 |
| | Planctomycetes_unclassified | 0,084 | 0,072 |
| | BD7-11 | 0,038 | 0,103 |
| | Pla3_lineage | 0,025 | 0,027 |

| | | | |
|---|---|---|---|
| | 028H05-P-BN-P5 | 0,003 | 0,005 |
| | ODP123 | 0,002 | 0,005 |
| Poribacteria | Poribacteria_cl | 0,009 | 0,010 |
| Proteobacteria | | | |
| | Gammaproteobacteria | 19,718 | 10,596 |
| | Deltaproteobacteria | 7,736 | 3,122 |
| | Alphaproteobacteria | 6,577 | 6,946 |
| | Proteobacteria_unclassified | 0,293 | 0,133 |
| | Magnetococcia | 0,010 | 0,071 |
| Rokubacteria | NC10 | 0,640 | 0,532 |
| Schekmanbacteria | Schekmanbacteria_cl | 0,019 | 0,020 |
| Spirochaetes | | 34,335 | 13,574 |
| | MVP-15 | 0,080 | 0,139 |
| | uncultured | 0,068 | 0,074 |
| | Spirochaetia | 0,021 | 0,027 |
| | Leptospirae | 0,010 | 0,022 |
| | Spirochaetes_cl | 0,001 | 0,003 |
| TA06 | TA06_cl | 0,006 | 0,010 |
| Tenericutes | Mollicutes | 0,119 | 0,781 |
| Verrucomicrobia | Verrucomicrobiae | 0,729 | 0,380 |
| WOR-1 | WOR-1_cl | 0,084 | 0,059 |
| WPS-2 | WPS-2_cl | 0,033 | 0,056 |
| WS2 | WS2_cl | 0,011 | 0,013 |
| WS4 | WS4_cl | 0,016 | 0,019 |
| Zixibacteria | Zixibacteria_cl | 0,158 | 0,082 |

Table S4. Relative abundance (%) of each bacterial class and phylum in the groundwater of H52.

| Phylum | Class | Mean (%) | Standard deviation (%) |
|---|---|---|---|
| Acidobacteria | | 0.455 | 0.230 |
| | Acidobacteriia | 0.209 | 0.150 |
| | Subgroup_6 | 0.100 | 0.065 |
| | Holophagae | 0.076 | 0.057 |
| | Thermoanaerobaculia | 0.033 | 0.039 |
| | Subgroup_17 | 0.020 | 0.021 |
| | Blastocatellia_(Subgroup_4) | 0.009 | 0.017 |
| | AT-s3-28 | 0.008 | 0.014 |
| Actinobacteria | | 2.836 | 1.876 |
| | Actinobacteria | 1.745 | 1.563 |
| | Acidimicrobiia | 0.771 | 0.451 |
| | Thermoleophilia | 0.265 | 0.236 |
| | Actinobacteria_unclassified | 0.036 | 0.043 |
| | Coriobacteriia | 0.011 | 0.016 |
| | WCHB1-81 | 0.005 | 0.008 |
| | RBG-16-55-12 | 0.002 | 0.004 |
| Armatimonadetes | | 0.018 | 0.046 |
| | uncultured | 0.012 | 0.045 |
| | Fimbriimonadia | 0.007 | 0.008 |
| Bacteria_unclassified | Bacteria_unclassified | 5.159 | 1.589 |
| Bacteroidetes | | 5.666 | 2.999 |
| Bacteroidetes | Ignavibacteria | 3.305 | 2.001 |
| | Bacteroidia | 2.341 | 1.396 |
| | Bacteroidetes_unclassified | 0.020 | 0.023 |
| BHI80-139 | BHI80-139_cl | 0.006 | 0.008 |
| BRC1 | BRC1_cl | 0.002 | 0.004 |
| Chlamydiae | Chlamydiae | 0.012 | 0.018 |
| Chloroflexi | | 0.886 | 0.402 |
| | Anaerolineae | 0.300 | 0.188 |
| | JG30-KF-CM66 | 0.223 | 0.184 |
| | Dehalococcoidia | 0.155 | 0.125 |
| | KD4-96 | 0.128 | 0.099 |
| | Chloroflexi_unclassified | 0.027 | 0.025 |
| | Chloroflexia | 0.027 | 0.031 |
| | OLB14 | 0.019 | 0.018 |
| | P2-11E | 0.006 | 0.008 |
| Cyanobacteria | | 0.067 | 0.052 |
| | Sericytochromatia | 0.051 | 0.045 |
| | Melainabacteria | 0.015 | 0.027 |
| | Cyanobacteria_unclassified | 0.001 | 0.002 |
| Dependentiae | Babeliae | 0.089 | 0.129 |
| Elusimicrobia | | 1.350 | 0.697 |
| | Elusimicrobia | 1.152 | 0.580 |
| | Lineage_IIc | 0.185 | 0.146 |

| | | | |
|---|---|---|---|
| | Elusimicrobia_cl | 0.009 | 0.015 |
| | Elusimicrobia_unclassified | 0.003 | 0.006 |
| Epsilonbacteraeota | Campylobacteria | 0.215 | 1.428 |
| FCPU426 | FCPU426_cl | 0.001 | 0.003 |
| Fibrobacteres | Fibrobacteria | 0.013 | 0.014 |
| Firmicutes | Clostridia | 0.178 | 0.261 |
| GAL15 | GAL15_cl | 0.004 | 0.006 |
| Gemmatimonadetes | Gemmatimonadetes | 0.837 | 0.503 |
| Hydrogenedentes | Hydrogenedentia | 0.037 | 0.046 |
| Kiritimatiellaeota | Kiritimatiellae | 0.012 | 0.031 |
| Latescibacteria | Latescibacteria_cl | 0.087 | 0.066 |
| Lentisphaerae | | 0.141 | 0.160 |
| | Lentisphaeria | 0.119 | 0.153 |
| | Oligosphaeria | 0.023 | 0.035 |
| Lindowbacteria | Lindowbacteria_cl | 0.004 | 0.009 |
| Margulisbacteria | Margulisbacteria_cl | 0.106 | 0.085 |
| Nitrospinae | | 0.030 | 0.077 |
| | Nitrospinia | 0.026 | 0.077 |
| | P9X2b3D02 | 0.004 | 0.011 |
| Nitrospirae | | 12.117 | 4.607 |
| | Thermodesulfovibrionia | 11.769 | 4.613 |
| | Nitrospira | 0.196 | 0.161 |
| | 4-29-1 | 0.138 | 0.204 |
| | HDB-SIOI1093 | 0.007 | 0.010 |
| | Nitrospirae_unclassified | 0.007 | 0.014 |
| Omnitrophicaeota | | 1.759 | 1.504 |
| | Omnitrophicaeota_cl | 1.741 | 1.503 |
| | Omnitrophia | 0.018 | 0.017 |
| Patescibacteria | | 35.008 | 10.734 |
| | Parcubacteria | 22.858 | 9.263 |
| | ABY1 | 9.221 | 4.656 |
| | Gracilibacteria | 1.436 | 1.043 |
| | Saccharimonadia | 0.859 | 0.673 |
| | Patescibacteria_unclassified | 0.471 | 0.278 |
| | Berkelbacteria | 0.093 | 0.101 |
| | Microgenomatia | 0.063 | 0.088 |
| | CPR2 | 0.007 | 0.009 |
| | WWE3 | 0.001 | 0.003 |
| PAUC34f | PAUC34f_cl | 0.009 | 0.014 |
| Planctomycetes | | 7.222 | 3.669 |
| | Brocadiae | 6.849 | 3.561 |
| | Phycisphaerae | 0.223 | 0.176 |
| | OM190 | 0.049 | 0.035 |
| | Planctomycetacia | 0.025 | 0.020 |
| | Pla4_lineage | 0.023 | 0.034 |
| | Planctomycetes_unclassified | 0.019 | 0.023 |

| | | | |
|---|---|---|---|
| | Pla3_lineage | 0.013 | 0.016 |
| | BD7-11 | 0.011 | 0.018 |
| | vadinHA49 | 0.009 | 0.013 |
| Proteobacteria | | 25.123 | 9.987 |
| | Gammaproteobacteria | 17.561 | 8.647 |
| | Deltaproteobacteria | 6.131 | 2.379 |
| | Alphaproteobacteria | 1.066 | 0.643 |
| | Proteobacteria_unclassified | 0.365 | 0.193 |
| Rokubacteria | NC10 | 0.057 | 0.048 |
| Spirochaetes | | 0.076 | 0.082 |
| | MVP-15 | 0.051 | 0.081 |
| | Leptospirae | 0.016 | 0.019 |
| | Spirochaetia | 0.009 | 0.011 |
| Tenericutes | Mollicutes | 0.001 | 0.004 |
| Verrucomicrobia | Verrucomicrobiae | 0.261 | 0.179 |
| WOR-1 | WOR-1_cl | 0.063 | 0.058 |
| WPS-2 | WPS-2_cl | 0.012 | 0.021 |
| Zixibacteria | Zixibacteria_cl | 0.081 | 0.051 |

Table S5. The effect of groundwater recharge on bacterial 16S rRNA gene abundance over time in groundwater wells. Since the values were not normally distributed (based on Shapiro-Wilk test of normality), we used non-parametric methods to test the effect of period and recharge. The effect of Period was tested using one-factorial Kruskal Wallis rank sum test.

| **Well** | Mean | SD | **Period** | Mean | SD | **Recharge** | Mean | SD |
|---|---|---|---|---|---|---|---|---|
| ***H41*** | 9.30E+07 | 4.20E+08 | ***P13*** | 9.1E+05 | 5.3E+05 | ***Recharge*** | 9.10E+05 | 5.30E+05 |
| | | | ***P15*** | 1.7E+07 | 1.2E+07 | ***Recession*** | 1.70E+07 | 1.20E+07 |
| | | | | | | ***Recharge*** | 1.40E+07 | 1.30E+07 |
| | | | ***P16*** | 1.4E+07 | 8.6E+06 | ***Recession*** | 1.60E+07 | 9.40E+06 |
| | | | | | | ***Recharge*** | 1.10E+07 | 7.70E+06 |
| | | | ***P17*** | 3.5E+07 | 1.5E+07 | ***Recession*** | 3.90E+07 | 1.60E+07 |
| | | | | | | ***Recharge*** | 2.80E+07 | 1.40E+07 |
| | | | ***P18*** | 1.7E+08 | 1.7E+08 | ***Recession*** | 2.50E+07 | 7.30E+06 |
| | | | | | | ***Recharge*** | 2.60E+08 | 7.50E+07 |
| | | | ***P19*** | 3.0E+08 | 9.1E+08 | ***Recession*** | 8.80E+07 | 2.90E+07 |
| | | | | | | ***Recharge*** | 6.30E+08 | 1.30E+09 |
| ***H43*** | 3.20E+07 | 3.70E+07 | ***P13*** | 2.4E+06 | 1.1E+06 | ***Recharge*** | 2.40E+06 | 1.10E+06 |
| | | | ***P14*** | 2.4E+07 | 1.8E+07 | ***Recession*** | 4.10E+06 | 6.20E+06 |
| | | | | | | ***Recharge*** | 3.40E+07 | 1.20E+07 |
| | | | ***P15*** | 2.6E+07 | 3.1E+07 | ***Recession*** | 2.30E+07 | 2.70E+07 |
| | | | | | | ***Recharge*** | 2.90E+07 | 3.70E+07 |
| | | | ***P16*** | 8.4E+06 | 5.2E+06 | ***Recession*** | 5.30E+06 | 2.50E+06 |
| | | | | | | ***Recharge*** | 1.20E+07 | 5.20E+06 |
| | | | ***P17*** | 4.9E+07 | 6.1E+07 | ***Recession*** | 2.00E+07 | 1.10E+07 |
| | | | | | | ***Recharge*** | 1.10E+08 | 8.00E+07 |
| | | | ***P18*** | 5.3E+07 | 3.6E+07 | ***Recession*** | 6.60E+07 | 5.00E+07 |
| | | | | | | ***Recharge*** | 4.80E+07 | 3.30E+07 |
| | | | ***P19*** | 4.2E+07 | 3.2E+07 | ***Recession*** | 2.20E+07 | 9.20E+06 |
| | | | | | | ***Recharge*** | 6.60E+07 | 3.30E+07 |
| ***H52*** | 3.20E+08 | 2.30E+08 | ***P13*** | 3.3E+07 | 6.5E+06 | ***Recharge*** | 3.30E+07 | 6.50E+06 |
| | | | ***P15*** | 1.8E+08 | 1.1E+08 | ***Recession*** | 1.70E+08 | 1.10E+08 |
| | | | | | | ***Recharge*** | 2.00E+08 | 1.20E+08 |
| | | | ***P16*** | 1.9E+08 | 8.1E+07 | ***Recession*** | 2.10E+08 | 7.80E+07 |
| | | | | | | ***Recharge*** | 1.70E+08 | 8.60E+07 |
| | | | ***P17*** | 4.3E+08 | 1.4E+08 | ***Recession*** | 4.30E+08 | 1.50E+08 |
| | | | | | | ***Recharge*** | 4.10E+08 | 7.90E+07 |
| | | | ***P18*** | 3.9E+08 | 1.5E+08 | ***Recession*** | 4.50E+08 | 1.70E+08 |
| | | | | | | ***Recharge*** | 3.30E+08 | 1.10E+08 |
| | | | ***P19*** | 5.9E+08 | 3.2E+08 | ***Recession*** | 7.60E+08 | 2.60E+08 |
| | | | | | | ***Recharge*** | 3.70E+08 | 2.80E+08 |

Table S6. The effect of groundwater recharge on bacterial alpha diversity (Shannon index) over time in groundwater wells.

| **Well** | Mean | SD | **Period** | Mean | SD | **Recharge** | Mean | SD |
|---|---|---|---|---|---|---|---|---|
| ***H41*** | 6.7 | 0.8 | ***P13*** | 6.2 | 0.2 | ***Recharge*** | 6.2 | 0.2 |
| | | | ***P15*** | 6.7 | 0.6 | ***Recession*** | 6.7 | 0.7 |
| | | | | | | ***Recharge*** | 6.7 | 0.4 |
| | | | ***P16*** | 6.6 | 0.6 | ***Recession*** | 6.3 | 0.2 |
| | | | | | | ***Recharge*** | 7.0 | 0.8 |
| | | | ***P17*** | 6.8 | 0.4 | ***Recession*** | 6.6 | 0.3 |
| | | | | | | ***Recharge*** | 7.2 | 0.5 |
| | | | ***P18*** | 8.0 | 0.7 | ***Recession*** | 7.4 | 0.5 |
| | | | | | | ***Recharge*** | 8.4 | 0.5 |
| | | | ***P19*** | 6.2 | 1.0 | ***Recession*** | 5.7 | 0.5 |
| | | | | | | ***Recharge*** | 6.5 | 1.3 |
| ***H43*** | 7.2 | 1.0 | ***P13*** | 6.2 | 0.6 | ***Recharge*** | 6.2 | 0.6 |
| | | | ***P14*** | 7.0 | 0.7 | ***Recession*** | 6.7 | 1.2 |
| | | | | | | ***Recharge*** | 7.2 | 0.4 |
| | | | ***P15*** | 6.9 | 1.0 | ***Recession*** | 7.3 | 0.7 |
| | | | | | | ***Recharge*** | 6.3 | 1.2 |
| | | | ***P16*** | 7.4 | 0.5 | ***Recession*** | 7.2 | 0.4 |
| | | | | | | ***Recharge*** | 7.7 | 0.5 |
| | | | ***P17*** | 7.7 | 0.8 | ***Recession*** | 7.5 | 0.7 |
| | | | | | | ***Recharge*** | 8.3 | 0.8 |
| | | | ***P18*** | 7.9 | 1.7 | ***Recession*** | 9.4 | 0.1 |
| | | | | | | ***Recharge*** | 7.4 | 1.7 |
| | | | ***P19*** | 6.9 | 0.5 | ***Recession*** | 7.1 | 0.2 |
| | | | | | | ***Recharge*** | 6.8 | 0.7 |
| ***H52*** | 5.4 | 0.5 | ***P13*** | 5.4 | 0.0 | ***Recharge*** | 5.4 | 0.0 |
| | | | ***P15*** | 5.5 | 0.7 | ***Recession*** | 5.5 | 0.7 |
| | | | | | | ***Recharge*** | 5.2 | 0.7 |
| | | | ***P16*** | 5.2 | 0.4 | ***Recession*** | 5.4 | 0.4 |
| | | | | | | ***Recharge*** | 5.1 | 0.4 |
| | | | ***P17*** | 5.0 | 0.3 | ***Recession*** | 5.0 | 0.3 |
| | | | | | | ***Recharge*** | 4.9 | 0.4 |
| | | | ***P18*** | 5.7 | 0.5 | ***Recession*** | 5.7 | 0.8 |
| | | | | | | ***Recharge*** | 5.6 | 0.3 |
| | | | ***P19*** | 5.7 | 0.2 | ***Recession*** | 5.8 | 0.3 |
| | | | | | | ***Recharge*** | 5.7 | 0.2 |

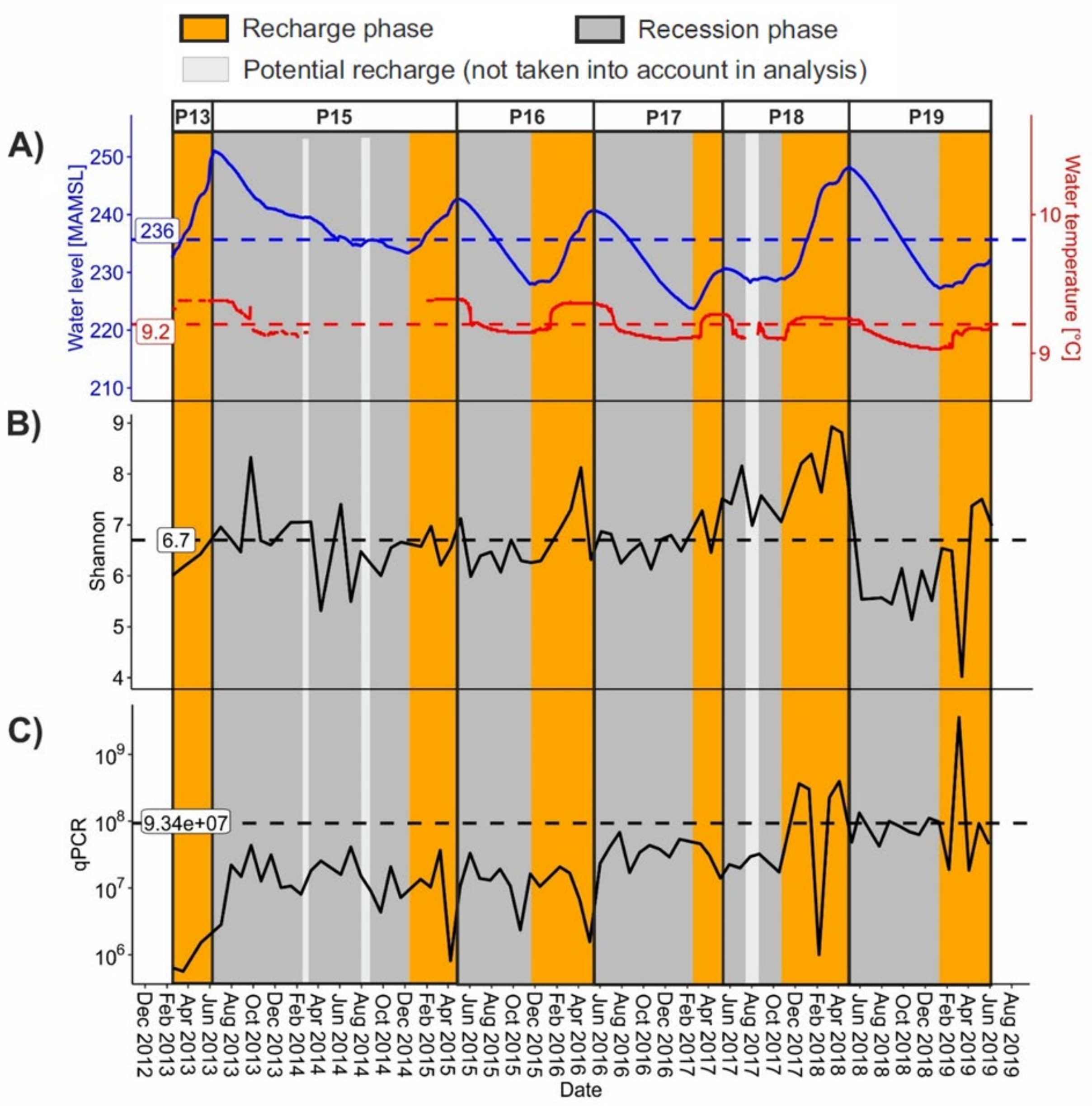


Figure S1. Dynamic changes of A) groundwater level and temperature on a daily basis, B) alpha diversity and C) bacterial 16S rRNA gene copy numbers (qPCR) on a 4-week basis in H41 well. "Period" is defined for hypothesis testing based on water level changes (increase: recharge; decrease: recession). A period (P) includes one recharge and one recession phase. P0 has no non-recharge phase.

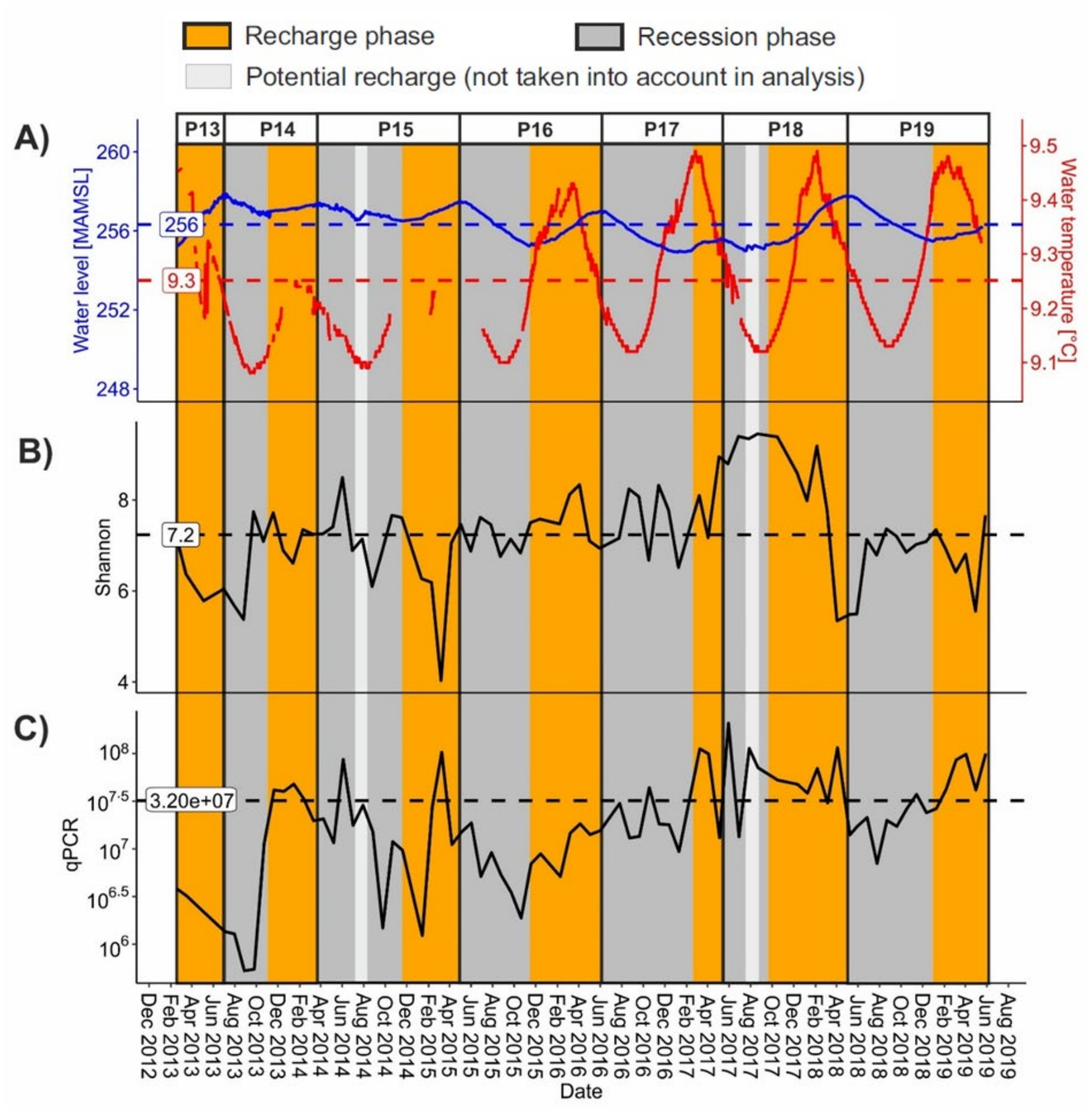


Figure S2. Dynamic changes of A) groundwater level and temperature on a daily basis, B) alpha diversity and C) bacterial 16S rRNA gene copy numbers (qPCR) on a 4-week basis in H43 well. "Period" is defined for hypothesis testing based on water level changes (increase: recharge; decrease: recession). A period (P) includes one recharge and one recession phase. P0 has no non-recharge phase.

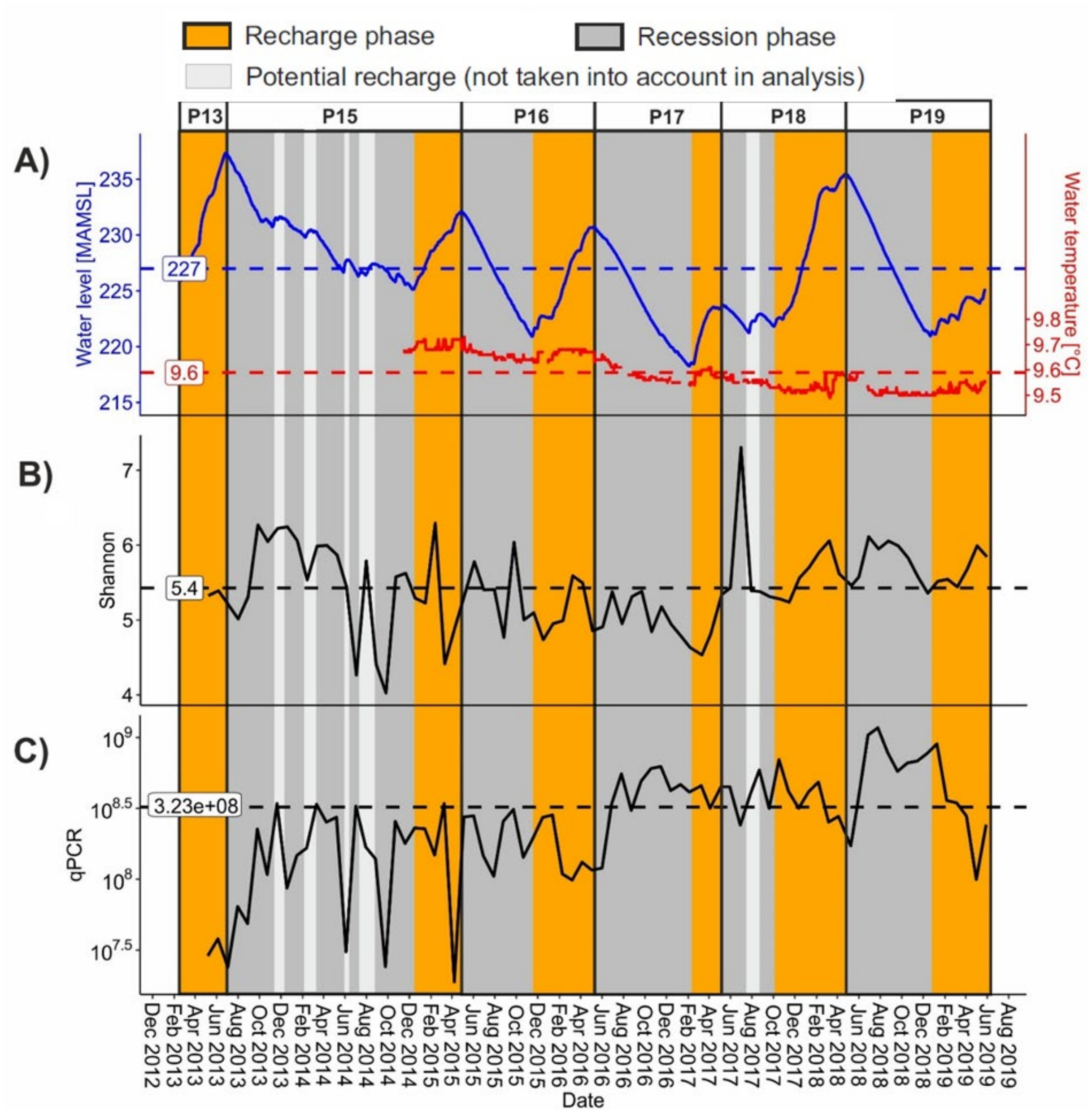


Figure S3. Dynamic changes of A) groundwater level and temperature on a daily basis, B) alpha diversity and C) bacterial 16S rRNA gene copy numbers (qPCR) on a 4-week basis in H52 well. "Period" is defined for hypothesis testing based on water level changes (increase: recharge; decrease: recession). A period (P) includes one recharge and one recession phase. P0 has no non-recharge phase.

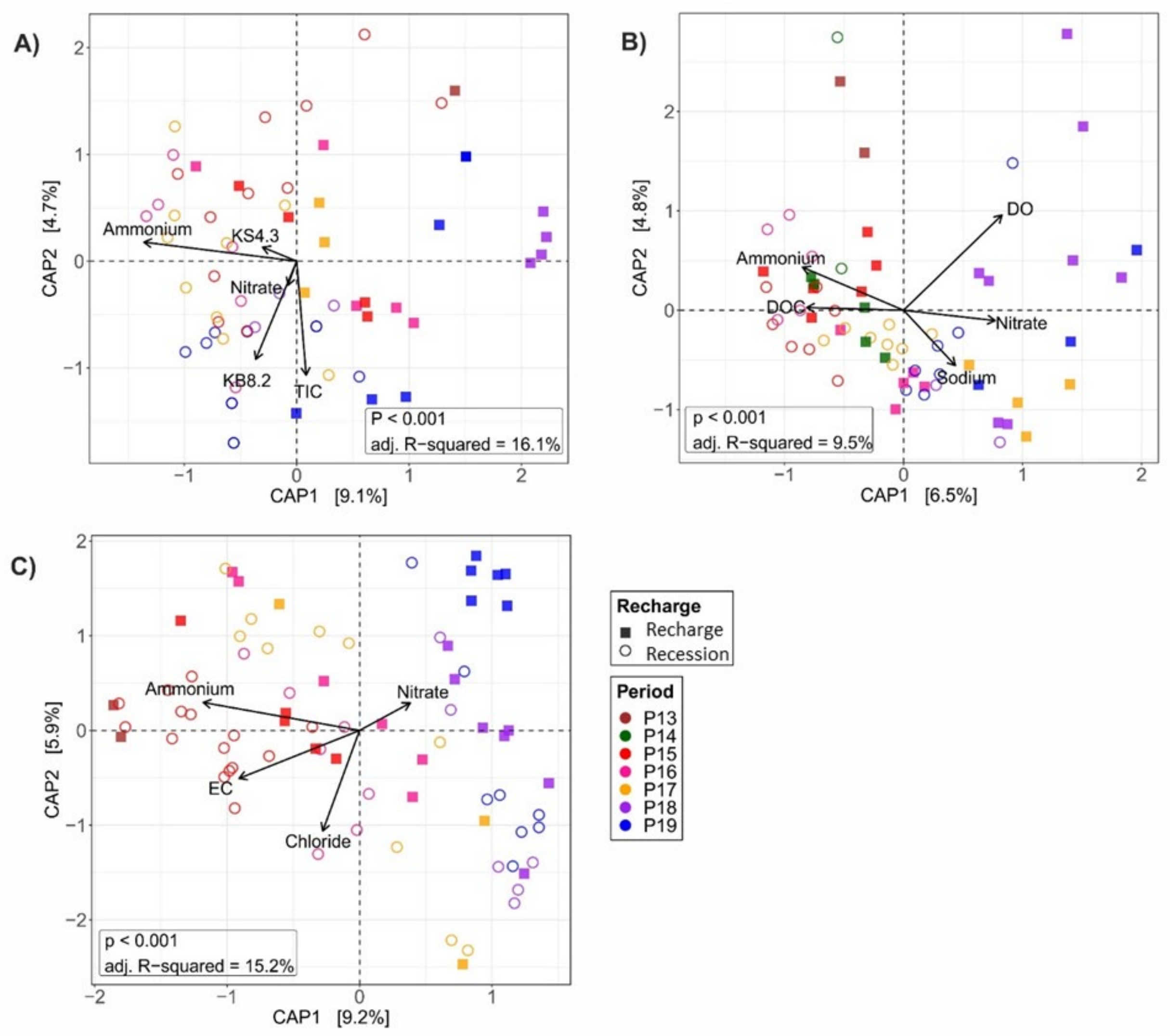


Figure S4. Distance-based RDA plots displaying the relationships between bacterial communities and significant environmental variables in the groundwater wells of A) H41, B) H43 and C) H52. Only significant environmental factors ($p < 0.05$) were plotted as vectors in the db-RDA plots based on 999 permutations. The symbols in the brackets following the vector labels indicate the significant effect of the constraining hydrochemical variables on the community data (*** $p < 0.001$, ** $p < 0.01$, * $p < 0.05$). The symbols in the brackets following the vector labels indicate the significant effect of the constraining hydrochemical variables on the community data (*** $p < 0.001$, ** $p < 0.01$, * $p < 0.05$). Abbreviations: *DO* dissolved oxygen, *DOC* dissolved organic carbon, *EC* electrical conductivity (specific), *TIC* total inorganic carbon, *KS4.3* acid neutralizing capacity (alkalinity), *KB8.2* base neutralizing capacity (acidity), *adj. R–squared* proportion of variance in bacterial community data explained by the hydrochemical parameters, adjusted for the number of predictors in the model.

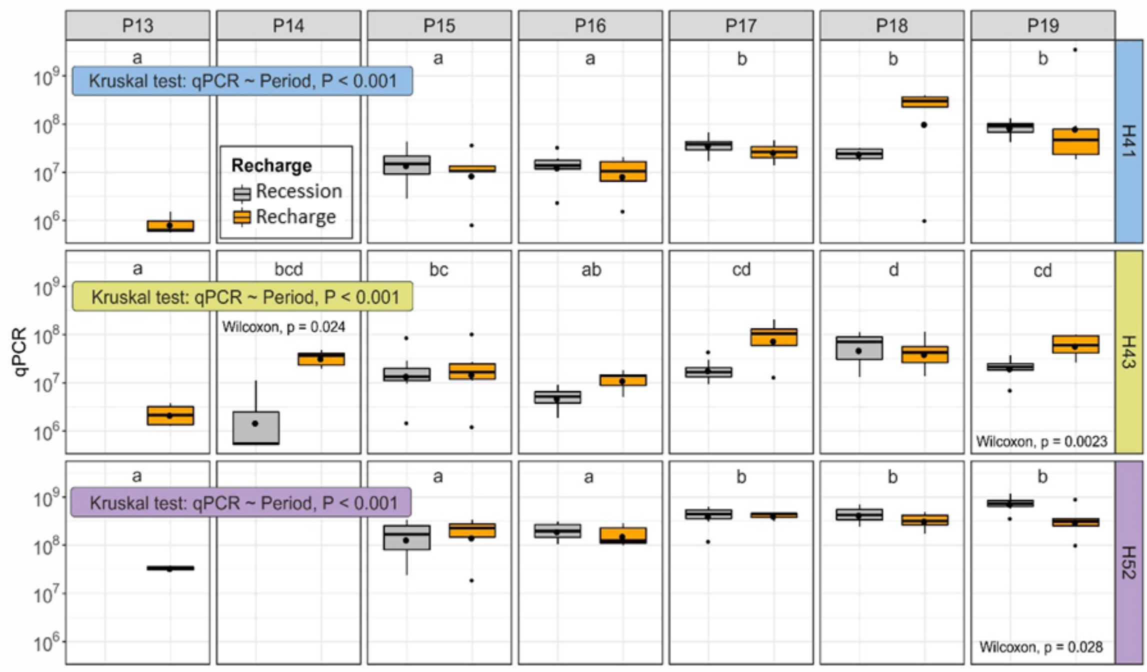


Figure S5. The effect of groundwater recharge on bacterial 16S rRNA gene abundance over time in groundwater wells. Since the values were not normally distributed (based on Shapiro-Wilk test of normality), we used non-parametric methods to test the effect of period and recharge. The effect of Period was tested using one-factorial Kruskal Wallis rank sum test. Different lowercase letters represent significant difference in 16S rRNA gene copy numbers between the periods based on Dunn test for multiple comparisons. The effect of recharge was tested using unpaired two-samples Wilcoxon test within each period.

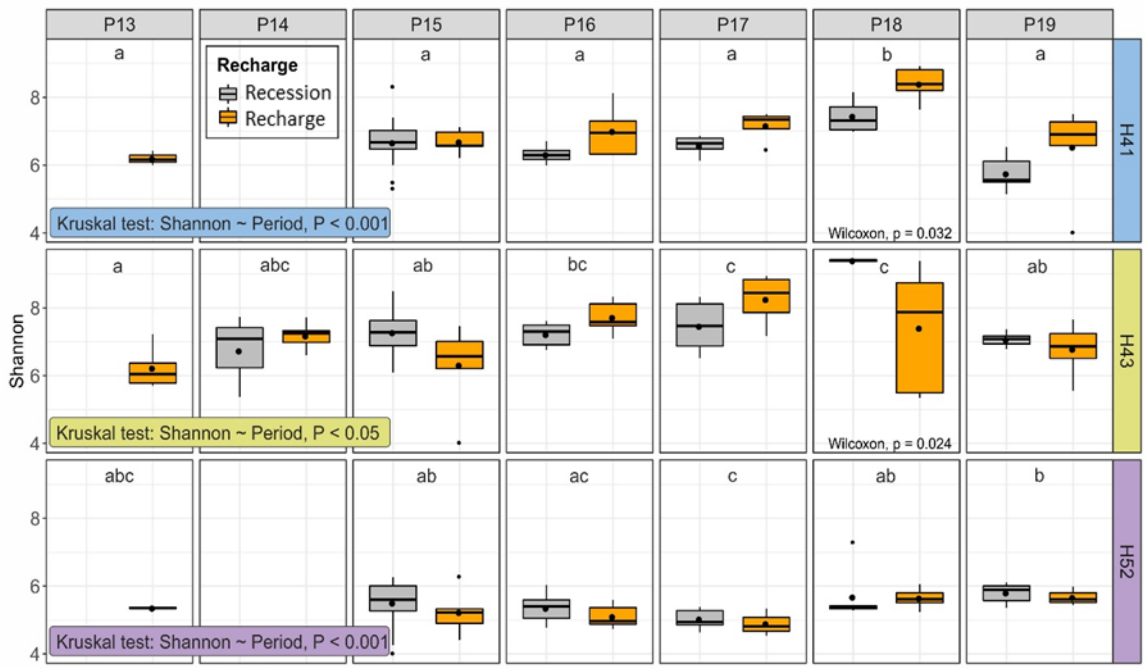


Figure S6. The effect of groundwater recharge on bacterial alpha diversity (Shannon index) over time in groundwater wells. Since the values were not normally distributed (based on Shapiro-Wilk test of normality), we used non-parametric methods to test the effect of period and recharge. The effect of Period was tested using one-factorial Kruskal Wallis rank sum test. Different lowercase letters represent significant difference in Shannon diversity values between the periods based on Dunn test for multiple comparisons. The effect of recharge was tested using unpaired two-samples Wilcoxon test within each period.

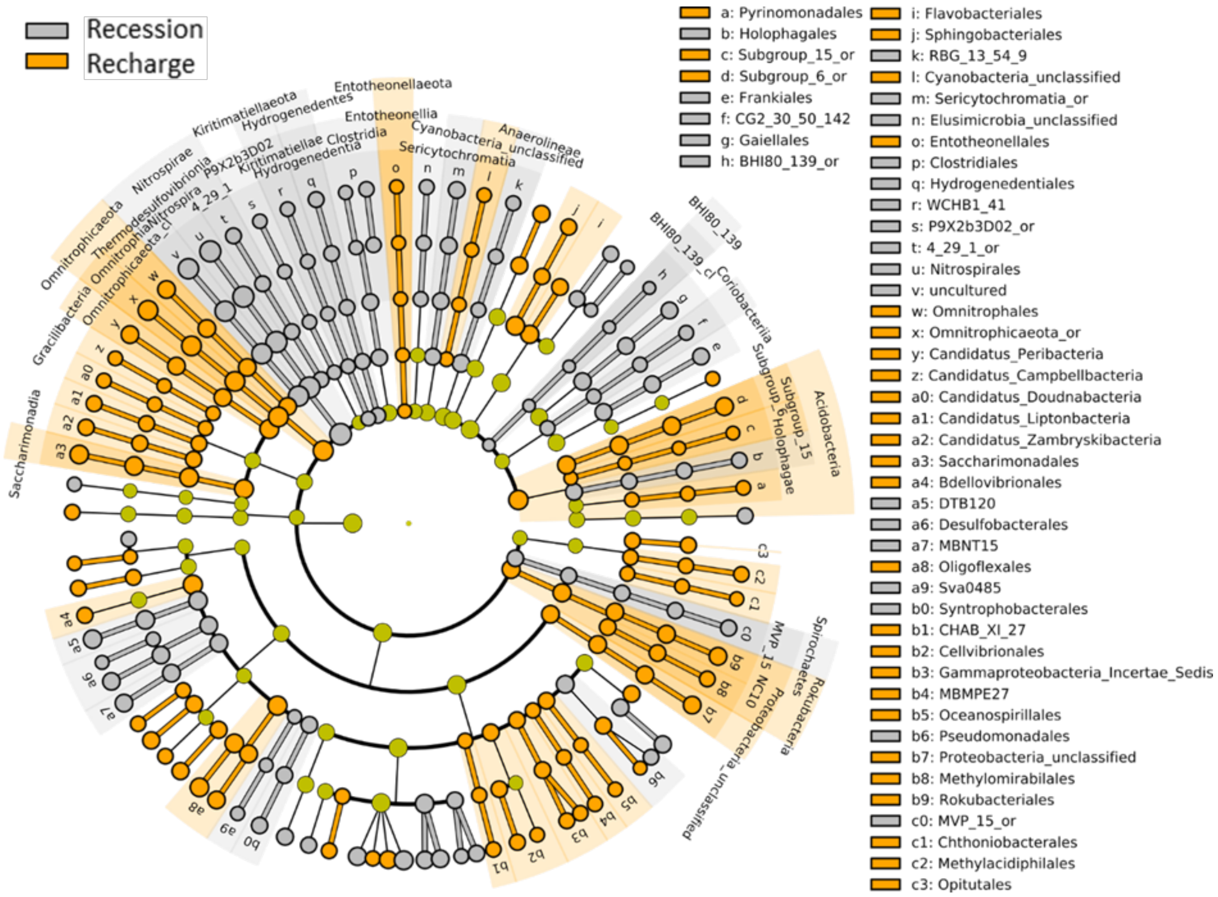


Figure S7. The discriminative features of bacterial taxa between recharge and recession phases in H41 well, tested based on LEfSe method ($p < 0.05$, LDA effect size > 2). For clarity, the cladogram presented here only displays the discriminative features of bacterial taxa at the phylum, class and order level.

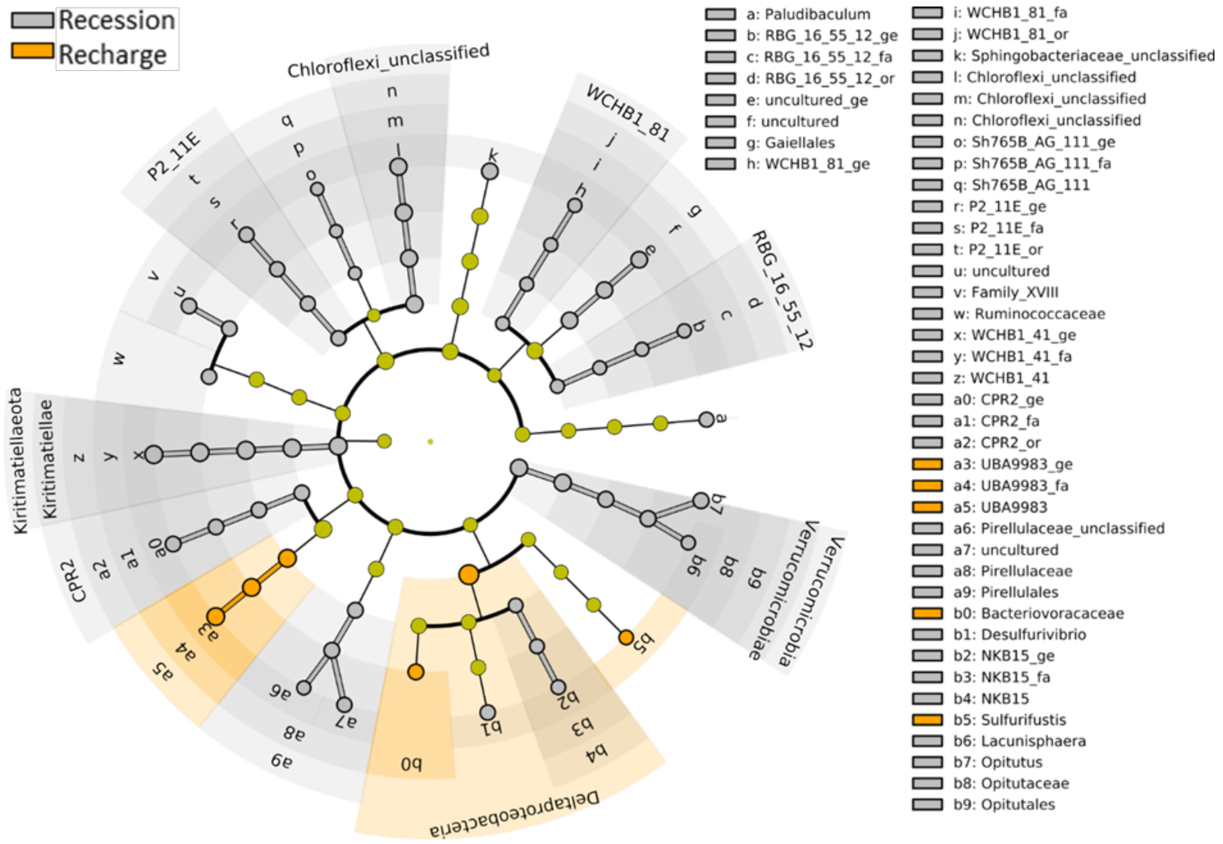


Figure S8. A cladogram showing the discriminative features of bacterial taxa between recharge and recession phases in H43 well at the phylum, class, order, family and genus level, tested with LEfSe method ($p < 0.05$, LDA effect size > 2).

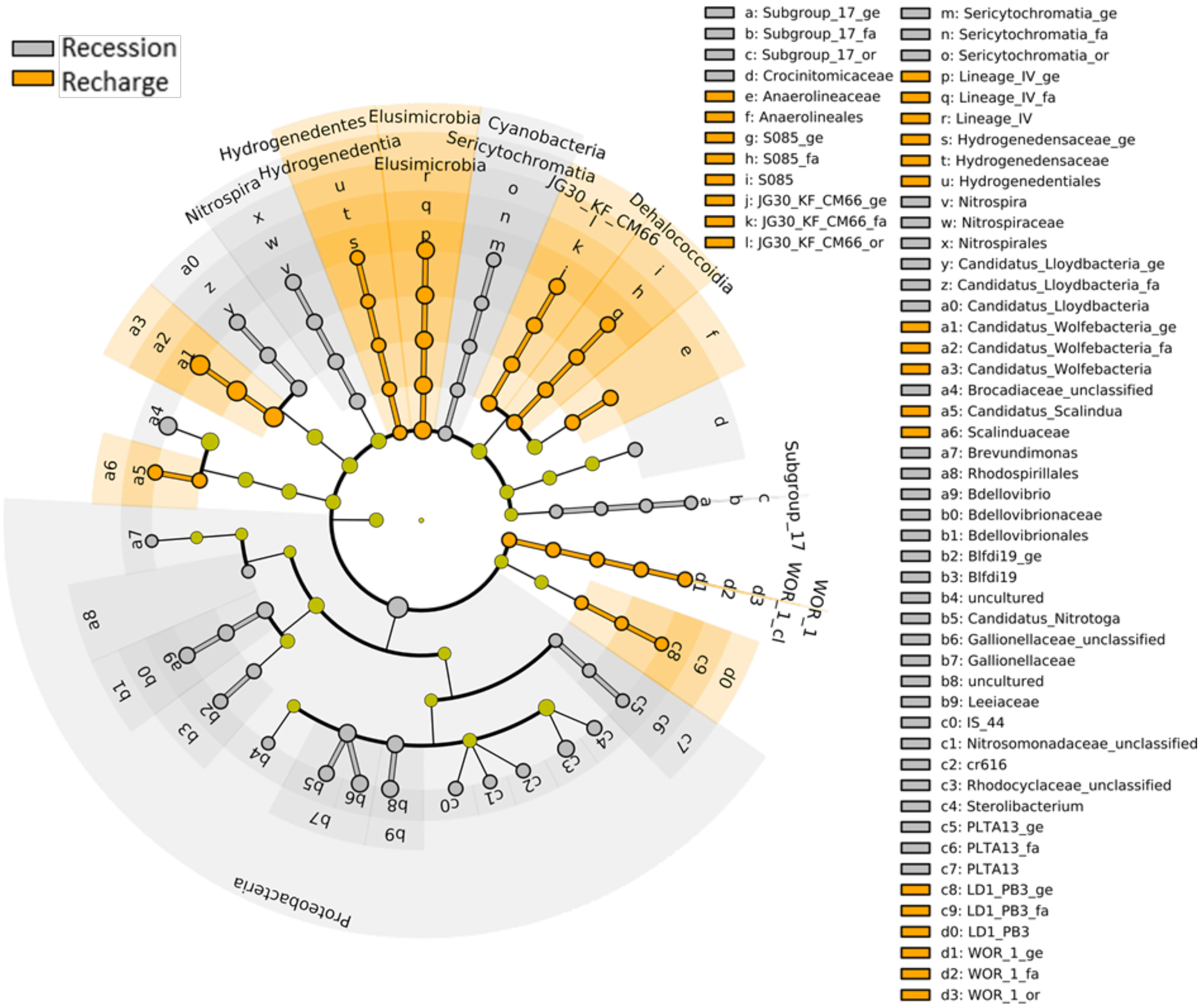


Figure S9. A cladogram showing the discriminative features of bacterial taxa between recharge and recession phases in H52 well at the phylum, class, order, family and genus level level, tested with LEfSe method ($p < 0.05$, LDA effect size > 2).
